# APP β-CTF triggers cell-autonomous synaptic toxicity independent of Aβ

**DOI:** 10.1101/2024.07.11.603028

**Authors:** Mengxun Luo, Jia Zhou, Cailu Sun, Wanjia Chen, Chaoying Fu, Chenfang Si, Yaoyang Zhang, Yang Geng, Yelin Chen

## Abstract

Aβ is believed to play a significant role in synaptic degeneration observed in Alzheimer’s disease (AD) and is primarily investigated as a secreted peptide. However, the contribution of intracellular Aβ or other cleavage products of its precursor protein (APP) to synaptic loss remains uncertain. In this study, we conducted a systematic examination of their cell-autonomous impact using a sparse expression system. Here, these proteins/peptides were overexpressed in a single neuron, surrounded by thousands of untransfected neurons. Surprisingly, we found that APP induced dendritic spine loss only when co-expressed with BACE1. This effect was mediated by β-CTF, a β-cleavage product of APP, through an endosome-related pathway independent of Aβ. Neuronal expression of β-CTF in mouse brains resulted in defective synaptic transmission and cognitive impairments, even in the absence of amyloid plaques. These findings unveil a β-CTF-initiated mechanism driving synaptic toxicity irrespective of amyloid plaque formation and suggest a potential intervention by inhibiting the endosomal GTPase Rab5.

## Introduction

Alzheimer’s disease (AD) stands as the most prevalent form of neurodegenerative disease and a leading cause of dementia. It is characterized by extracellular amyloid plaque deposition, intracellular neurofibrillary tangles, synaptic and neuronal loss, and neuroinflammation (Blennow et al., 2006; Gómez-Isla et al., 1996; JohnHardy and Selkoe, 2002; Leyns and Holtzman, 2017; Selkoe, 2002). The amyloid plaque primarily consists of aggregated Aβ, a cleavage product of amyloid precursor protein (APP) (Glenner and Wong, 1984; Masters et al., 1985). Notably, two Aβ antibodies have demonstrated efficacy in removing amyloid plaques and slowing AD progression in phase III clinical trials (Sims et al., 2023; van Dyck et al., 2023). Furthermore, numerous naturally occurring mutations in genes encoding APP or its catalytic enzyme γ-secretase result in early onset familial AD or reduce AD risk (Jonsson et al., 2012; Liu et al., 2017; Mullan et al., 1992; Nilsberth et al., 2001). This collective evidence, spanning AD pathology, human genetics, and intervention trials, strongly supports a causal role of Aβ and amyloid plaque in AD pathogenesis. However, despite clinical trials employing Aβ antibodies targeting Aβ oligomers, protofibrils, or deposited plaque, AD progression has been slowed down by only ∼30% (Mintun et al., 2021; Pleen and Townley, 2022; Sims et al., 2023; van Dyck et al., 2023). Notably, antibodies against monomeric soluble Aβ failed to yield clinical benefits (Sperling et al., 2023). It is possible that these Aβ antibodies may overlook certain pathogenic factors crucial for AD pathogenesis.

APP, a type I transmembrane protein, undergoes cleavage primarily by α-secretase on the cytoplasmic membrane, producing soluble α-cleavage N-terminal fragment (sAPPα) and α-cleavage C-terminal fragment (α-CTF). Some APP molecules bypass α-cleavage and undergo endocytosis into endocytic compartments, where they are subsequently cleaved by β-secretase, generating soluble β-cleavage N-terminal fragment (sAPPβ) and β-cleavage C-terminal fragment (β-CTF). β-CTF is further cleaved by γ-secretase to produce Aβ and APP intracellular domain (AICD) (Golde et al., 1992; Zhang and Song, 2013; Zhang et al., 2011). While extracellular Aβ, targeted by Aβ antibodies, is widely studied, the potential contribution of intracellular Aβ, APP, and other APP cleavage products to AD pathogenesis remains uncertain (Konietzko, 2012; Kwart et al., 2019; Nikolaev et al., 2009; Oddo et al., 2003; Vohra et al., 2010; Willem et al., 2015). For instance, β-CTF has been implicated in endosomal dysfunction(Israel et al., 2012; Jiang et al., 2010; Kim et al., 2016; Kwart et al., 2019; Xu et al., 2016), yet its downstream functional impacts remain unclear.

Synapse loss represents an early feature of AD neurodegeneration and is closely associated with cognitive dysfunction (de Wilde et al., 2016; DeKosky et al., 1996; Terry et al., 1991). Secreted Aβ induces synaptic dysfunction by interacting with its receptors on the neuronal plasma membrane (Kamenetz et al., 2003; Kessels et al., 2013; Wei et al., 2010). Other studies have reported γ-secretase inhibition reduced spine density *in vivo* via an APP-dependent pathway (Bittner et al., 2009). Additionally, deletion of APP in mice has been shown to decrease dendritic spine density (Tyan et al., 2012). The diverse outcomes of APP on synapse suggest a complex impact of APP and metabolites. It remains unclear whether APP or other APP fragments can also induce synaptic toxicity in a cell-autonomous manner.

To address these inquiries, we employed a sparse transfection system utilizing the Helios gene gun to explore the potential role of intracellular Aβ or other APP fragments. These molecules were expressed in a single neuron surrounded by untransfected wild type neurons. Surprisingly, full-length APP did not induce synaptic toxicity. However, co-expression of APP with BACE1 resulted in significant loss of dendritic spines, indicating a crucial role of APP β-cleavage in synaptic damage. Further investigations unveiled that this detrimental effect was mediated by β-CTF in a cell-autonomous manner, independent of Aβ. Additionally, *in vivo* expression of β-CTF was adequate to induce synaptic dysfunction and cognitive impairments in mice, even in the absence of amyloid plaques. In summary, our study delineates a mechanism initiated by β-CTF that can induce synaptic degeneration in a cell-autonomous manner, thus extending beyond the scope of Aβ antibody-based therapies.

## Results

### APP only led to spine loss when co-expressed with BACE1

To investigate the cell-autonomous impact of APP on neurons, we utilized a Helios gene gun transfection system to sparsely express APP in rat organotypic hippocampal slice cultures (Chen et al., 2014). In hippocampal slice cultures at DIV9, transient expression of a familial AD APP mutation (APP_Swedish_) (Mullan et al., 1992) with GFP in CA1 pyramidal neurons for six days did not reduce their dendritic spine densities compared with neurons expressing GFP alone (Figure 1A-B), suggesting that APP alone does not significantly contribute to synaptic loss.

**Figure 1.**
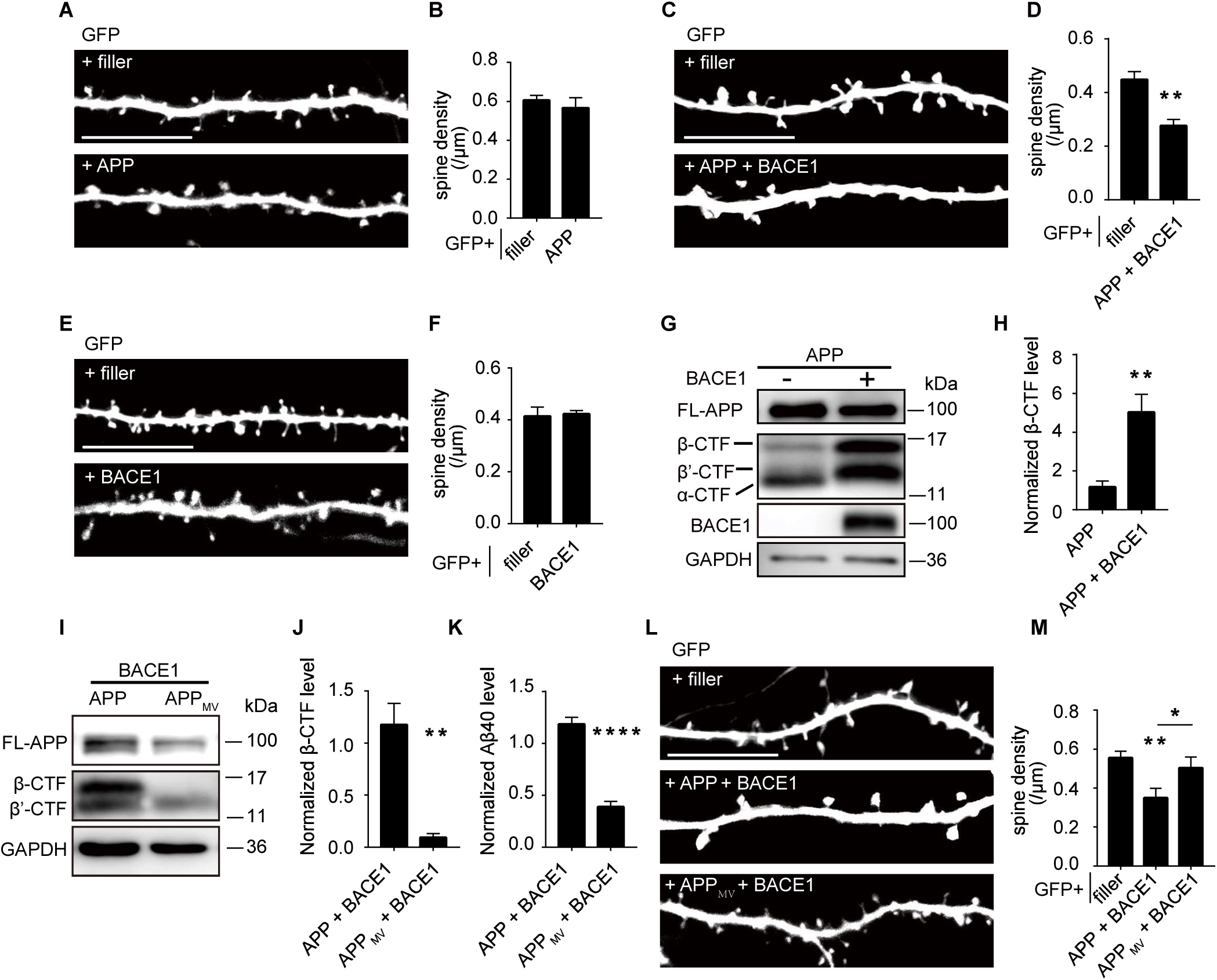
APP only led to spine loss when co-expressed with BACE1. **A-B** Representative images and spine density of basal dendrites from CA1 pyramidal neurons transiently expressing GFP alone or together with APP. (filler, n =21; APP, n=16). **C-D** Representative images and spine density of basal dendrites from CA1 pyramidal neurons transiently expressing GFP alone or together with APP and BACE1 with a ratio of 15:1. (filler, n=8; APP plus BACE1, n=5) **E-F** Representative images and spine density of basal dendrites from CA1 pyramidal neurons transiently expressing GFP alone or together with BACE1. (filler, n=15; BACE1, n=15) **G** Western blot and corresponding statistical analysis of APP (Y188), BACE1 and GAPDH from HEK293T cells expressing APP or APP plus BACE1 with a ratio of 15:1. GAPDH was measured as a loading control. **H** Measurement of APP β-CTF (β C-Terminal Fragment) level from HEK293T cells expressing APP alone or APP and BACE1 with a ratio of 15:1. n=4. **I** Western blot and corresponding statistical analysis of APP fragments (Y188) from HEK293T cells expressing APP plus BACE1 or APP_MV_ plus BACE1. **J** Measurement of APP β-CTF levels from HEK293T cells expressing APP plus BACE1 or APP_MV_ plus BACE1. n=4. **K** Measurement of Aβ40 levels from HEK293T cells expressing APP plus BACE1 or APP_MV_ plus BACE1. n=4. **L-M** Representative images and spine density of basal dendrites from CA1 pyramidal neurons transiently expressing GFP alone or together with APP and BACE1 or APP_MV_ and BACE1. (filler, n=12; APP plus BACE1, n=14; APP_MV_ plus BACE1, n=10) All dendritic images were acquired from rat organotypic hippocampal slice cultures after transfection for 6-7 days. Statistics: One-way ANOVA or Student’s t test. * p < 0.05, ** p < 0.01, *** p < 0.001, **** p <0.0001. Error bars show SEM. Scale bars, 10 μm.

BACE1-mediated β-cleavage of APP is crucial for amyloidogenesis, and BACE1 activity is notably elevated in AD patient brains (Cheng et al., 2014; Vassar et al., 1999). In human brains, APP and BACE1 are expressed at a ratio of about 15:1 (Uhlén et al., 2015). Endogenous BACE1 levels may not suffice to cleave the overexpressed APP. To assess whether insufficient β-cleavage of APP underlies the lack of synaptic toxicity caused by overexpressed APP, we co-expressed APP with BACE1 (in a 15:1 ratio) in organotypic hippocampal slice cultures and observed a significant ∼40% reduction in spine density compared to neurons expressing GFP alone (Figure 1C-D). Transient expression of BACE1 alone did not affect spine density (Figure 1E-F). Co-expression of APP and BACE1 with different ratios or using an internal ribosome entry site (IRES) also resulted in significant spine loss (Supplementary figure 1A-B). These findings support the requirement of BACE1 for APP to induce synaptic loss.

When expressed alone in HEK293T cells, APP is predominantly cleaved at its α site to produce sAPPα and α-CTF (C-terminal fragment after APP α-cleavage) (Figure 1G, Supplementary figure 1C). Co-expression with BACE1 led to increased cleavage of APP at its β and β’ sites, resulting in elevated β-CTF (more than 4-fold) and β’-CTF as the major metabolic products and a substantial reduction in α-CTF levels (Figure 1G-H, Supplementary figure 1C). APP_MV_ (M596V) cannot be cleaved by BACE1 to produce β-CTF and Aβ but has no impact on β’-cleavage (Figure 1I-K) (Citron et al., 1995). When co-expressed with BACE1, APP_MV_ failed to induce spine loss (Figure 1L-M), supporting the requirement of β-cleavage of APP to induce spine loss.

In summary, these findings suggest that certain β-cleavage products of APP, rather than APP or BACE1 alone, could lead to spine loss in a cell-autonomous manner.

### β-CTF induced spine loss independent of Aβ

APP undergoes cleavage by secretases, generating soluble N-terminal fragments (sAPPs) and C-terminal fragment (CTFs). We then explored the impact of different sAPPs and APP-CTFs on dendritic spines. Transient expression of β-CTF significantly reduced the spine density of CA1 pyramidal neurons in organotypic rat hippocampal cultures, whereas expression of sAPPα, sAPPβ, α-CTF or β’-CTF did not produce such an effect (Figure 2A-B, Supplementary figure 2A-B). APP CTFs exhibited similar expression levels in HEK293T cells (Figure 2C). When the plasmid amount was reduced to 1/8 of the original dose, β-CTF no longer induced a decrease in dendritic spine density (Supplementary figure 2E-F), indicating the synaptic damaging effect of β-CTF is its expression level dependent.

**Figure 2.**
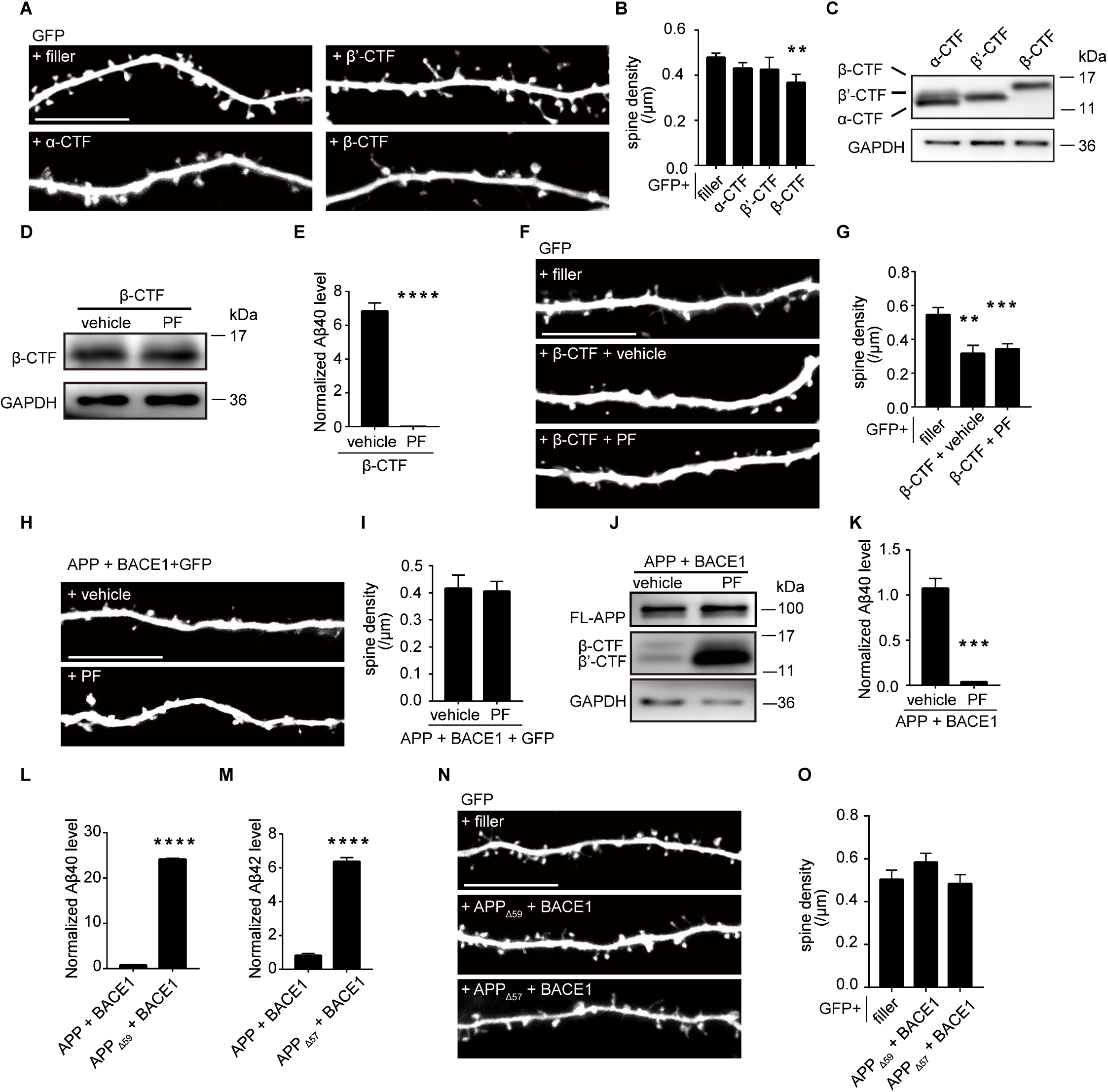
β-CTF induced spine loss independent of Aβ. **A-B** Representative images and spine density of basal dendrites from CA1 pyramidal neurons transiently expressing GFP alone or together with α-CTF, β’-CTF or β-CTF.(filler, n=19; α-CTF, n=7; β’-CTF, n=8; β-CTF, n=7) **C** Western blot of APP fragments from HEK293T cells expressing α-CTF-flag, β’-CTF-flag and β-CTF-flag (flag antibody). **D** Western blot of APP fragments from HEK293T cells expressing β-CTF-flag treatment with vehicle or PF (flag antibody). **E** Measurement of Aβ40 secreted from HEK293T cells expressing β-CTF treatment with vehicle or PF. n=4. **F-G** Representative images and spine density of basal dendrites from CA1 pyramidal neurons transiently expressing GFP alone or together with β-CTF after treated with vehicle or PF. (filler, n=11; β-CTF, n=7, β-CTF plus PF, n=14) **H-I** Representative images and spine density of basal dendrites from CA1 pyramidal neurons transiently expressing GFP together with APP and BACE1 after treated with vehicle or PF. (APP plus BACE1 with vehicle, n=9; APP plus BACE1 with PF, n=14) **J** Western blot of APP fragments (Y188) from HEK293T cells co-expressing APP and BACE1 after treated with vehicle or PF. **K** Measurements of Aβ40 from HEK 293T cells expressing APP and BACE1 after treated with vehicle or PF. n=4. **L** Measurements of Aβ40 from HEK293T cells expressing APP and BACE1 or APP_Δ59_ and BACE1. n=4. **M** Measurements of Aβ42 from HEK293T cells expressing APP and BACE1 or APP_Δ57_ and BACE1. n=4. **N-O** Representative images and spine density of basal dendrites from CA1 pyramidal neurons transiently expressing GFP alone or together with APP_Δ59/Δ57_ and BACE1. (filler, n=9; APP_Δ59_ and BACE1, n=7; APP_Δ57_ and BACE1, n=6) All dendritic images were acquired from rat organotypic hippocampal slice cultures after transfection for 6-7 days. PF, PF03084014, a γ secretase inhibitor. Statistics: One-way ANOVA or Student’s t test. * p < 0.05, ** p < 0.01, *** p < 0.001, **** p <0.0001. Error bars show SEM. Scale bars, 10 μm.

β-CTF undergoes processing by γ-secretase to generate Aβ and AICD (APP intracellular domain) (Thinakaran and Koo, 2008). Subsequently, we investigated whether Aβ generated from β-CTF was responsible for the β-CTF-induced spine loss. Treatment with a γ-secretase inhibitor, PF03084014 (1μM), effectively reduced Aβ to baseline levels in cells expressing β-CTF without altering the expression levels of β-CTF itself (Figure 2D-E). PF03084014 treatment did not affect the spine density in neurons expressing GFP alone (Supplementary figure 2C-D). Notably, PF03084014 treatment failed to prevent the spine loss induced by β-CTF expression (Figure 2F-G), or by co-expression of APP and BACE1 (Figure 2H-K), suggesting that Aβ might not be the causative factor.

To delve deeper into the impact of Aβ on dendritic spines, we engineered two APP mutants (APP_Δ59_ and APP_Δ57_) lacking the AICD domain, which are known to generate significant amounts of Aβ40 and Aβ42, respectively (Figure 2L-M). Co-expression of either APP_Δ59_ or APP_Δ57_ with BACE1, did not alter spine density (Figure 2N-O), further bolstering the idea that Aβ is not responsible for the β-CTF-induced spine loss. Subsequently, we investigated the involvement of another β-CTF cleavage product, AICD, which has been reported to interact with the transcription factor forkhead box O (FoxO) and promote FoxO-induced transcription of proapoptotic genes, leading to cell death (Wang et al., 2014). However, transient expression of AICD failed to alter the density of spines (Supplementary figure 2G-H).

In conclusion, these results support the idea that APP can induce spine loss in a cell-autonomous manner through β-CTF, independent of Aβ and AICD.

### Expression of β-CTF damaged synapses in mice

To ascertain whether β-CTF-induced spine loss could manifest *in vivo*, we examined the density of dendritic spines from CA1 pyramidal neurons infected with lentivirus encoding β-CTF and GFP, or GFP alone, in adult mice. Sparse expression of GFP and β-CTF was observed in CA1 pyramidal neurons (Figure 3A). The spine density of CA1 pyramidal neurons expressing β-CTF and GFP was significantly lower than that of neurons expressing GFP alone (Figure 3B-C), indicating that β-CTF could induce spine loss in a cell-autonomous manner *in vivo*.

**Figure 3.**
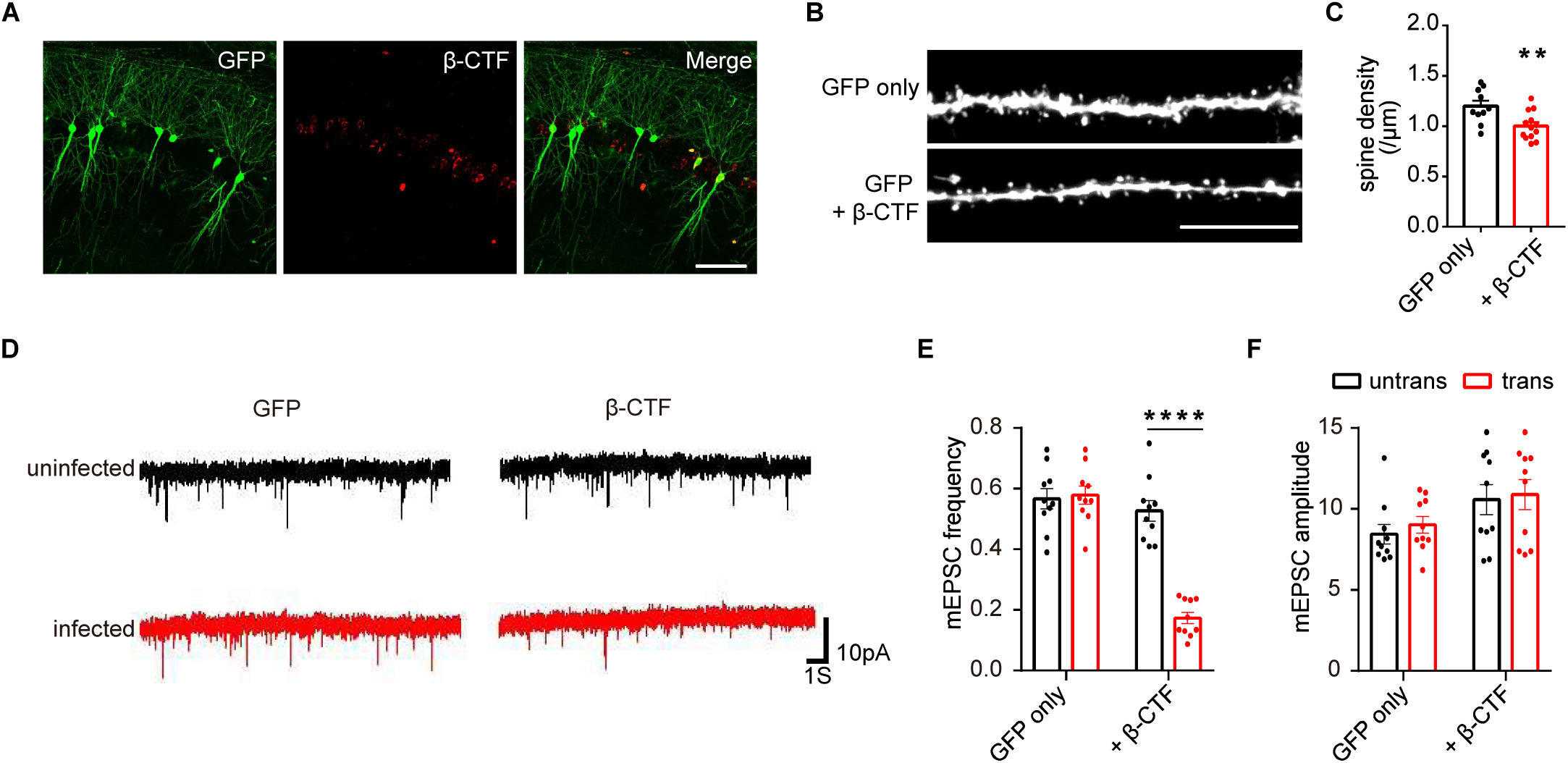
Expression of β-CTF damaged synapses in mouse brains. **A** Representative images of adult mouse CA1 pyramidal neurons infected with lentivirus expressing GFP and β-CTF. Scale bar, 100 μm. **B-C** Representative images from basal dendrites and quantitation of their spine density in CA1 pyramidal neurons infected with lentivirus expressing GFP alone or GFP with β-CTF in mouse hippocampi. Scale bar, 10 μm. GFP only, n=10; GFP plus β-CTF, n=13. **D-F** Representative recording traces and quantitations of mEPSCs from adult mouse hippocampal neurons infected with lentivirus (identified with GFP signal) and neighboring uninfected neurons. n=10. Statistics: Two-way ANOVA or Student’s t test. * p < 0.05, ** p < 0.01, *** p < 0.001, **** p <0.0001. Error bars show SEM.

Next, we evaluated excitatory synaptic transmission through whole-cell patch-clamp recording of hippocampal pyramidal neurons infected with lentivirus encoding β-CTF or GFP. Neurons expressing β-CTF exhibited a ∼65% lower frequency of mEPSCs compared to neighboring uninfected neurons, whereas there was no significant change in mEPSC frequency in neurons expressing GFP alone (Figure 3D-E). However, mEPSC amplitude remained unaltered in neurons expressing either β-CTF or GFP (Figure 3D, F). These findings collectively suggest that β-CTF leads to reduced excitatory synapse density without affecting the strength of the remaining synapses.

### Expression of β-CTF damaged cognitive function in mice in the absence of plaque formation

We proceeded to investigate whether β-CTF affects cognitive functions. Adeno-associated viruses (AAV) encoding GFP or β-CTF were bilaterally injected into the hippocampus of 1-month-old mice (Figure 4A-B). After 4 months of expression, animal behaviors were examined. Immunostaining revealed widespread expression of GFP or β-CTF throughout the entire hippocampus (Figure 4B), with no detectable amyloid plaque formation observed using ThS staining (Figure 4C). Side-by-side staining demonstrated robust amyloid plaque deposition in the brain of two-month-old 5XFAD mice (Figure 4D), validating the efficacy of the staining method. The body weight of mice expressing β-CTF in the hippocampus was approximately 17% lower than that of GFP controls (Figure 4E). In the Y-maze test, the mean alternations were significantly reduced in mice expressing β-CTF compared to GFP controls, suggesting abnormal working memory in mice expressing β-CTF (Figure 4F).

**Figure 4.**
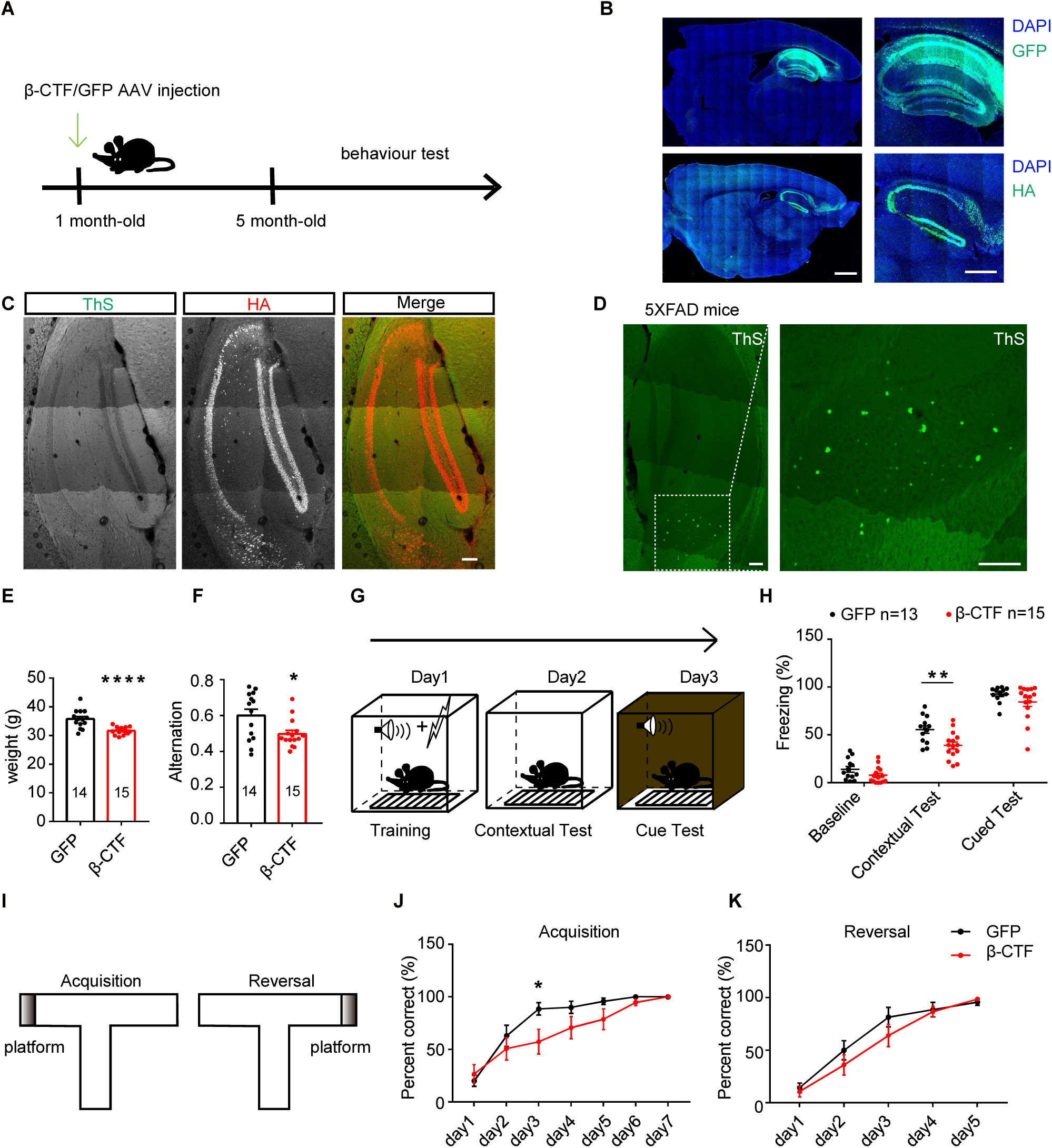
Expression of β-CTF damaged cognitive function in mice in the absence of plaque formation. **A** Schematic diagram showing the time line for stereotactic injection of AAV, behavioral training and tests. **B** Representative images showing immunofluorescence staining of expressed GFP or β-CTF-HA by AAV in mouse hippocampi (WT). Scare bars represent 1000 μm (left) and 500 μm (right). **C** Images of mouse hippocampus (WT) stained with ThS and HA antibody after infected with AAV encoding β-CTF-HA. Scale bar, 100 µm. **D** Representative image of ThS staining of a 5XFAD mice hippocampal slice. Scale bar, 100 μm. **E** Weight of mice infected with AAV encoding GFP or β-CTF in their hippocampi. GFP, n=14; β-CTF, n=15. **F** Quantitations of spontaneous alternation in Y-maze from mice infected with AAV encoding GFP or β-CTF. GFP, n=14; β-CTF, n=15. **G** Schematic diagram showing fear conditioning test design. **H** Quantitation of freezing in contextual and cued fear conditioning tests from mice infected with AAV encoding GFP and β-CTF in their hippocampi. GFP, n=13; β-CTF, n=15. **I** Schematic diagram showing water T maze test design. **J-K** The percentage of correct responses across the 5 trials of water T maze test was quantified in each day of Acquisition (J) or Reversal (K). GFP, n=14; β-CTF, n=15. Statistics: Repeated measures two-way ANOVA or Student’s t test. * p < 0.05, ** p < 0.01, *** p < 0.001, **** p <0.0001. Error bars show SEM.

The fear conditioning test assesses associative fear learning and memory (Figure 4G) (Xiao et al., 2018). In this test, mice expressing β-CTF exhibited similar baseline levels of freezing time as GFP controls (Figure 4H). However, in the hippocampus-dependent contextual fear conditioning test, mice expressing β-CTF showed a significantly shorter freezing time (approximately 40% less) than those expressing GFP after training (Figure 4H). In the amygdala-dependent cued fear conditioning test, these two groups performed similarly (Figure 4H). In the water T maze test (Figure 4I), mice expressing β-CTF displayed slower learning curves than GFP controls, indicating impairments in acquisition learning (Figure 4J) but not in reversal learning (Figure 4K).

In the open field test, mice expressing β-CTF traveled a longer distance (Supplementary figure 3A-C) and spent more time in the center area with a higher frequency of center entrances (Supplementary figure 3D-E) compared to mice expressing GFP. However, there were no differences in rearing frequency and duration between the two groups (Supplementary figure 3F-G). These results suggest that mice expressing β-CTF exhibited increased locomotion and reduced anxiety-like behaviors. Consistently, mice injected with AAV encoding β-CTF spent significantly more time exploring and traveled a greater distance in the open arms compared to the GFP controls in the EPM (Supplementary figure 3H-J). In tail suspension tests, these two groups exhibited similar levels of immobility (Supplementary figure 3K), indicating that β-CTF expression in the hippocampus did not alter depression-like behaviors.

In conclusion, these findings support that β-CTF expression is sufficient to disrupt hippocampus-dependent cognitive functions.

### The C-terminal YENPTY motif was necessary for β-CTF to induce endosomal dysfunction and synapse loss

Endosome abnormalities mediated by APP β-CTFs have been reported across various cell types, including human iPSC-induced neurons, PC12M cells, and N2a cells (Kim et al., 2016; Kwart et al., 2019; Xu et al., 2016). Next, we investigated whether β-CTF impacted endosomes in hippocampal neurons. Dissociated cultured hippocampal neurons were co-transfected with APP-CTFs and Rab5-GFP, an endosomal marker fused with GFP, at DIV7. In neurons expressing either Rab5-GFP alone or Rab5-GFP with α-CTF, Rab5 puncta appeared uniform and smoothly rounded (Figure 5A-B). However, in neurons co-transfected with β-CTF, Rab5 puncta were larger and exhibited less uniform shapes, often appearing lobular, and showed robust co-localization with β-CTF (Figure 5C, D). Notably, the morphology of lysosomes, as observed by Lamp1 staining, remained unaffected by the expression of α-or β-CTF in neurons (Supplementary figure 4A-D), thereby suggesting a specific interaction of β-CTF with endosomes.

**Figure 5.**
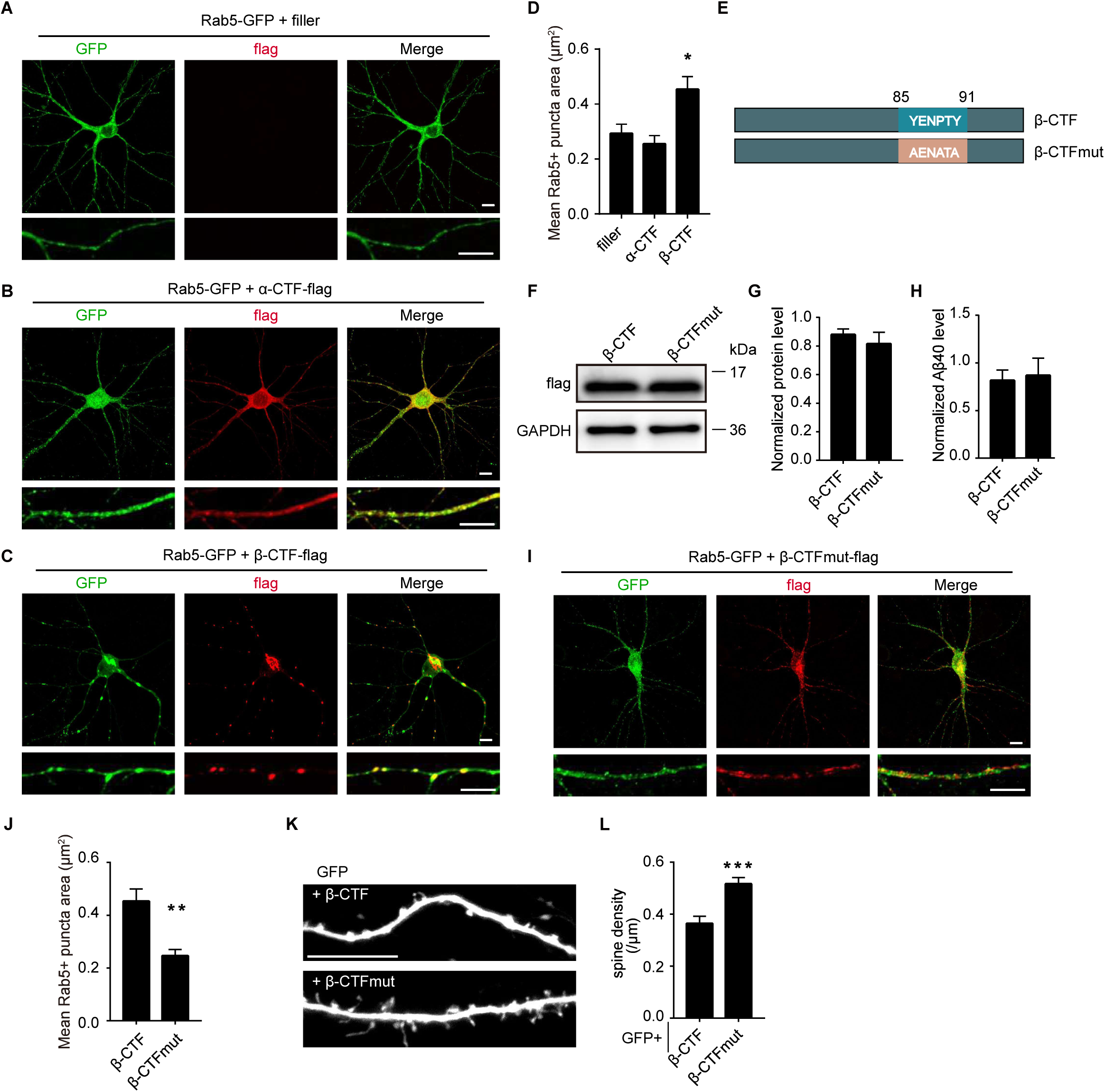
The C-terminal YENPTY motif was necessary for β-CTF to induce endosomal dysfunction and synapse loss. **A** Immunofluorescent staining of GFP (green) from dissociated rat hippocampal neurons expressing Rab5-GFP. **B** Immunofluorescent staining of GFP (green) and flag (red) from dissociated rat hippocampal neurons expressing Rab5-GFP and α-CTF-flag. **C** Immunofluorescent staining of GFP (green) and flag (red) from dissociated rat hippocampal neurons expressing Rab5-GFP and β-CTF-flag. **D** Quantitation of average Rab5+ puncta size of neurons expressing Rab5-GFP only or co-expressing α/β-CTF-flag and Rab5-GFP. (filler, n=9; α-CTF, n=9; β-CTF, n=11) **E** Schematic diagram of the β-CTF and β-CTFmut. **F** Western blot of APP fragments from HEK293T cells expressing β-CTF-flag and β-CTFmut-flag with a flag antibody. **G** Quantitations of APP fragments from HEK293T cells expressing β-CTF-flag and β-CTFmut-flag. n=4. **H** Measurement of Aβ40 from HEK 293T cells expressing β-CTF or β-CTFmut. n=5. **I** Immunofluorescent staining of GFP (green) and flag (red) from dissociated rat hippocampal neurons expressing Rab5-GFP and β-CTFmut-flag. **J** Quantitation of average Rab5+ puncta size of neurons co-expressing β-CTF-flag or β-CTFmut-flag and Rab5-GFP. (β-CTF, n=11; β-CTFmut, n=9) **K-L** Representative images and spine density of basal dendrites from CA1 pyramidal neurons transiently expressing GFP together with β-CTF or β-CTFmut for 6-7 days in rat organotypic hippocampal slice cultures. β-CTF, n=12; β-CTFmut, n=15. Scale bar, 10 μm. Statistics: One-way ANOVA or Student’s t test. * p < 0.05, ** p < 0.01, *** p < 0.001, **** p <0.0001. Error bars show SEM.

The C-terminal YENPTY motif of APP was found to be crucial for its interaction with endosomes (Lai et al., 1995). To elucidate the significance of this interaction, we expressed a mutant form of β-CTF (β-CTF_mut_), where the YENPTY motif was substituted with AENATA. Interestingly, β-CTF_mut_ was expressed at similar levels to wildtype β-CTF and exhibited comparable Aβ production (Figure 5E-H). However, unlike wildtype β-CTF, β-CTF_mut_ showed a more diffuse distribution throughout the neurons and failed to induce enlarged Rab5 puncta (Figure 5I-J). Notably, pyramidal neurons transiently expressing β-CTF_mut_ displayed a higher spine density compared to those expressing wildtype β-CTF (Figure 5K-L). These findings underscore the critical role of the YENPTY motif-mediated interaction with endosomes in β-CTF-induced spine loss.

### Spine loss induced by β-CTF was prevented by Rab5 inhibition

To investigate the downstream mechanism responsible for β-CTF-induced synaptic loss, we analyzed the proteomic alterations triggered by β-CTF in dissociated cultured hippocampal neurons (Figure 6A). Our findings revealed significant changes in protein levels upon exposure to β-CTF (Figure 6B). Notably, the expression of Synapsin-1, a presynaptic protein associated with synaptic vesicles, and GluR1, GluN2A, GluN2B, subunits of glutamate receptors, were diminished in neurons expressing β-CTF (Figure 6C-F). VAMP2, a key component of SNARE complex was also reduced by β-CTF (Figure 6E, G). The protein level of Synapsin-1 was further reduced after treatment with γ-secretase inhibitors in neurons expressing β-CTF (Supplementary figure 5A-B). Gene Ontology analysis further elucidated that β-CTF expression downregulated proteins involved in membrane trafficking, synaptic vesicle cycle, pre-synapse, and post-synapse (Figure 6H-I), aligning with previous observations indicating that β-CTF induces synaptic dysfunction and endosomal abnormalities(Zhou et al., 2019). Conversely, upregulated proteins were primarily associated with peptide metabolic processes and translation (Supplementary figure 5C-D).

**Figure 6.**
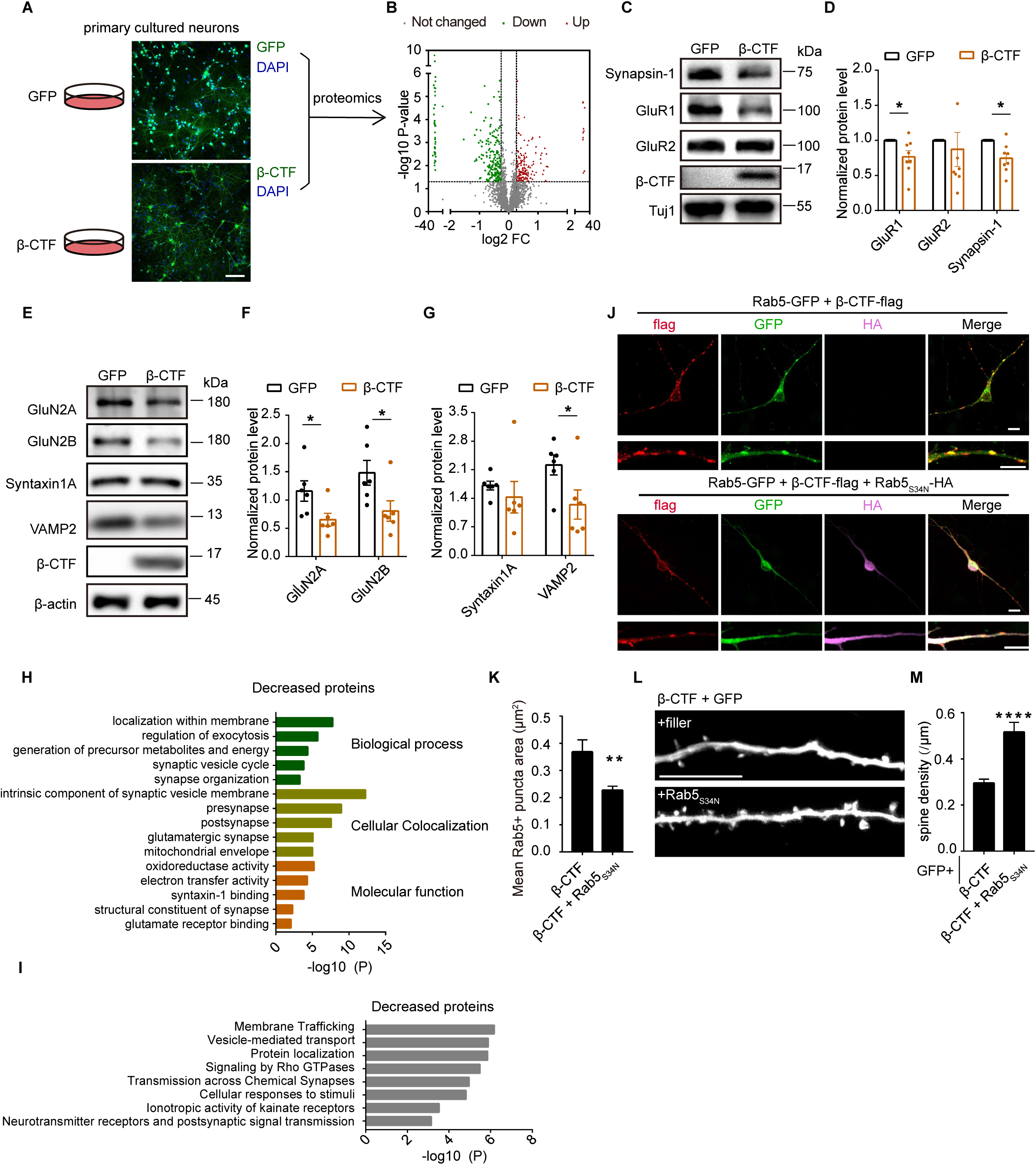
Spine loss induced by β-CTF was prevented by Rab5 inhibition. **A** Representative images of dissociated rat hippocampal neurons infected with lenti virus encoding GFP or β-CTF. Scale bar, 100 μm. **B** Volcano plot of quantitative mass spectrometry analysis showing protein omics changes in neurons infected with lenti virus encoding β-CTF v.s. GFP. Proteins with fold change >1.2 or <0.83 and p value < 0.05 are colored. **C-D** Western blot and corresponding statistic analysis of Synapsin-1, GluR1, GluR2 in dissociated hippocampal neurons infected with lentivirus expressing GFP or β-CTF. Tuj 1 was used as an internal control. **E-G** Western blot and corresponding statistic analysis of GluN2A, GluN2B, Syntaxin1A, VAMP2 in dissociated hippocampal neurons infected with lentivirus expressing GFP or β-CTF. β-actin was used as an internal control. **H** The Gene Ontology (GO) processes analysis of decreased proteins by Metascape. **I** The Reactome Gene Sets processes analysis of decreased protein by Metascape. **J** Immunofluorescent staining of GFP (green), flag (red) and HA (magenta) from dissociated rat hippocampal neurons expressing Rab5-GFP and β-CTF-flag or together with Rab5_S34N_-HA. Scale bar, 10 μm. **K** Quantitation of average Rab5+ puncta size from dissociated rat hippocampal neurons expressing Rab5-GFP and β-CTF-flag or together with Rab5_S34N_-HA. (β-CTF, n=5; β-CTF + Rab5_S34N_, n=9) **L-M** Representative images and measurements of spine density of basal dendrites from CA1 pyramidal neurons transiently expressing GFP and β-CTF with or without Rab5_S34N_ for 6-7 days in rat organotypic hippocampal slice cultures. Scale bar, 10 μm. (β-CTF, n=17; β-CTF plus Rab5_S34N_, n=11) Statistics: Student’s t test. * p < 0.05, ** p < 0.01, *** p < 0.001, **** p <0.0001. Error bars show SEM.

Overexpression of β-CTF resulted in the enlargement of Rab5-positive endosomes, similar to the effects observed with a constitutively active mutant of Rab5, Rab5_Q79L_ (Supplementary figure 5E-F) (Kim et al., 2016). Notably, the introduction of a dominant negative mutant of Rab5, Rab5_S34N_, attenuated the endosome enlargement induced by β-CTF (Figure 6J-K). In neurons expressing Rab5_S34N_, there was reduced co-localization of β-CTF with Rab5 (Figure 6J-K). Critically, co-expression of Rab5_S34N_ with β-CTF effectively mitigated the spine loss induced by β-CTF in hippocampal slice cultures (Figure 6L-M). These findings underscored that Rab5 overactivation-induced endosomal dysfunction contributed to β-CTF-induced spine loss. However, expression of Rab5_S34N_ in β-CTF-expressing neurons did not alter the levels of synapse-related proteins that were reduced in these neurons (Supplementary figure 5G-H), suggesting Rab5 overactivation did not contribute to these protein expression changes induced by β-CTF.

## Discussion

Treatment with Lecanemab or Donanemab, two Aβ antibodies, significantly slowed down AD progression by approximately 30% in phase III clinical trials (Sims et al., 2023; van Dyck et al., 2023). The success of these antibodies in modifying AD progression validates Aβ as a cause of AD pathogenesis. However, the relatively modest benefits have raised additional questions. Despite effectively reducing amyloid plaques to near baseline levels after 18 months of treatment in some patients, functional benefits were limited to only ∼30%. This raises the question of whether amyloid plaque is the primary source of Aβ-related toxicity or if these patients were treated too late to reverse their disease progression more effectively. This study aims to investigate whether Aβ-associated pathways could lead to pathogenesis beyond secreted Aβ and amyloid plaques, which falls outside the scope of Aβ antibody-based therapies due to their inability to penetrate cytoplasmic membranes.

AD is characterized by significant synaptic degeneration. Conventionally, Aβ is considered detrimental to synapses, thus synaptic loss in AD is attributed to various forms of Aβ, including amyloid plaques (Kamenetz et al., 2003; Oakley et al., 2006). However, conflicting studies have suggested that Aβ may actually promote synaptogenesis (Abramov et al., 2009; Bittner et al., 2009; Puzzo et al., 2008; Zhou et al., 2022). The exact role of Aβ in AD-related synaptic degeneration remains elusive. Most investigations have focused on the non-cell-autonomous function of Aβ after its secretion from neurons (Reinders et al., 2016). The potential contribution of intracellular Aβ or its precursors to synaptic toxicity has been largely unexplored. Utilizing a sparse neuron transfection system, we systematically examined this question. Intriguingly, among Aβ, APP, and major APP cleavage products, only β-CTF induced synaptic toxicity in a cell-autonomous manner (Figure 1, 2, Supplementary figure 1, 2), aligning with predictions based on indirect evidence (Kwart et al., 2019; Lee et al., 2022; Tamayev et al., 2012; Vaillant-Beuchot et al., 2021; Xu et al., 2016). In mouse brains, β-CTF also triggered substantial synaptic loss and cognitive deficits in the absence of amyloid plaques (Figure 4C), further supporting the notion that synaptic loss and amyloid plaque formation are mediated by distinct mechanisms. While current data did not exclude the potential involvement of Aβ-induced toxicity in the synaptic and cognitive dysfunction observed in mice overexpressing β-CTF, addressing this directly remains challenging. Treatment with γ-secretase inhibitors could potentially shed light on this issue. However, treatments with γ-secretase inhibitors are known to lead to brain dysfunction by itself likely due to its blockade of the γ-cleavage of other essential molecules, such as Notch (Doody et al., 2013; Güner and Lichtenthaler, 2020), preventing from pursuing it further experimentally *in vivo*.

APP, Aβ, and presenilins have been extensively studied in mouse models, providing convincing evidence that high Aβ concentrations are toxic to synapses (Chapman et al., 1999). Moreover, addition of Aβ to murine cultured neurons or brain slices is toxic to synapses (Wang et al., 2017). However, Aβ-induced synaptotoxicity was not observed in our study. A major difference between our study and others is that we employed an isolated expression system, where Aβ was applied solely to individual neurons surrounded by other neurons, without overwhelming them with excessive amounts of Aβ. In contrast, other studies typically apply Aβ to neurons indiscriminately. Therefore, we predict that Aβ does not lead to synaptic deficits from individual neurons in cell autonomous manners, whereas β-CTF does.

It is noteworthy that β-CTF induces spine loss independent of Aβ, implying that while synaptic degeneration and amyloidogenesis both occur downstream of β-CTF, they may represent two parallel processes independent of each other following APP β-cleavage. In fact, Aβ and β-CTF levels were significantly elevated in the AD brains (Kim et al., 2016). Aβ antibodies effectively facilitate the clearance of secreted Aβ, including that deposited in amyloid plaques. However, β-CTF localizes within the intracellular membrane of endo/lysosomal vesicles, rendering it inaccessible to Aβ antibodies. Consequently, Aβ antibody-based therapies cannot mitigate the neuronal toxicities initiated by β-CTF. This limitation could partially explain the restricted clinical benefits observed with Aβ antibodies. Addressing β-CTF-mediated synaptic toxicity should be prioritized in the future development of improved AD therapies, potentially in conjunction with Aβ antibodies.

However, clinical trials of BACE1 inhibitors have failed to demonstrate an efficacy in treating AD, despite their theoretical potential to inhibit the production of both β-CTF and Aβ. One potential reason is that pharmacological inhibition of BACE1 appears not as effective as its genetic removal. Genetic depletion of BACE1 leads to clearance of existing amyloid plaques (Hu et al., 2018), whereas pharmacological inhibition of BACE1 could not stop growth of existing plaques, although it slows down the growth of these plaques and prevents formation of new plaques (Peters et al., 2018). The development of better β-secretase inhibitors to more effectively inhibit β-cleavage is required to fully test the role of APP β-cleavage in AD pathogenesis.

Endosomal dysfunction is an early neuropathological signature of AD and Down syndrome (DS) mediated by β-CTF. Kim et al found β-CTF recruited APPL1 (a Rab5 effector) via YENPTY motif to Rab5 endosomes, where it stabilizes active GTP-Rab5, leading to pathologically accelerated endocytosis, endosome swelling and selectively impaired transport of Rab5 endosomes (Kim et al., 2016). Rab5 endosomal dysfunction may trigger prodromal and neurodegenerative features of Alzheimer’s disease (Pensalfini et al., 2020). However, the downstream functional impacts of endosomal dysfunction induced by β-CTF remain unclear. One of the impacts is deficits of NGF signaling in basal forebrain cholinergic neurons (BFCNs) (Salehi et al., 2006; Xu et al., 2016). Here, we report endosomal dysfunction-induced by β-CTF also contributes to synaptic dysfunction. Synaptic abnormalities are corrected by inhibiting Rab5 activity to restore endosomal function (Figure 6J-M), indicating that endosomal abnormalities may serve as a potential target for AD treatment.

β-CTF elicited significant spine loss, whereas α-CTF or β’-CTF, which are only 16 or 10 amino acids shorter at their N-terminal ends compared to β-CTF, did not exhibit this detrimental effect. Since the C-terminal YENPTY motif of β-CTF is essential for this activity, likely through its interaction with endosomal proteins, β-CTF represents the minimum size of the APP fragment capable of inducing cell-autonomous synaptic toxicity. However, the specific contribution of the N-terminal domain of β-CTF to synaptic loss remains unclear. It remains to be investigated whether longer fragments containing β-CTF could also induce synaptic loss. If confirmed, strategies aimed at reducing APP β-cleavage while promoting α-or β’-cleavage to decrease both Aβ and β-CTF may offer a more effective treatment for AD in the future.

## Materials and Methods

### Dissociated hippocampal neuron culture and transfection

We prepared hippocampal neurons from embryonic day 17/18 (E17/18) SD rats. Initially, whole hippocampi were dissected under stereomicroscopes and then digested into single cells using papain (Worthington) for 25 minutes at 37 ℃. The dissociated cells were seeded at a density of 180,000 cells per well in a 24-well plate, using Neurobasal media (Thermo Fisher Scientific) supplemented with 2% B27 (Life technology), 0.25% Glutamax (Thermo Fisher Scientific), and 100 units/mL Penicillin/Streptomycin (Life technology). Neurons were cultured in incubators maintained at 37 ℃ with 5% CO_2_, with fresh media half replenished every 5 days. For virus infection, dissociated hippocampal neurons at day *in vitro* (DIV) 14 were exposed to purified lentivirus. After 5 days, the infected neurons were lysed in RIPA buffer (Solarbio) supplemented with a protease inhibitor cocktail (Selleck) for further analysis. For plasmid transfection, dissociated hippocampal neurons at DIV7 were transfected with Lipofectamine 2000 (Thermo Fisher Scientific) following the manufacturer’s instructions.

### Western blotting and immunofluorescent staining

For western blotting, we first removed the culture medium, and then immediately placed neurons or HEK293T cells on ice. Subsequently, they were lysed using 2× loading buffer (100 mM Tris-HCl, 150 mM NaCl, 4% SDS, 20% glycerol). The lysates were agitated for 10 minutes at room temperature, followed by boiling at 98 ℃ for 10 minutes and centrifugation at 14,000 ×g for 10 minutes. The proteins were separated using 8%-12% Tris-glycine SDS-PAGE and transferred to a PVDF membrane (Millipore, 0.22 μm). Primary antibodies were then incubated in blocking buffer overnight at 4 °C, followed by secondary antibodies at room temperature for 2 hours. The signal was detected using an Amersham Imager 600 (GE Healthcare), and densitometry was measured using ImageJ.

For immunofluorescent staining, we first washed neurons on cover glasses with PBS (supplemented with 1 mM CaCl_2_ and 5 mM MgCl_2_) once. Then, we permeabilized them with 0.15% Triton-X in PBS for 15 minutes. After blocked with 5% bovine serum albumin in PBS for 30 minutes at room temperature, the samples were incubated with primary antibodies in blocking buffer at 4 °C overnight. The following day, the samples were washed three times with PBS and then incubated with secondary antibodies at room temperature for 1 hour in the dark. Subsequently, the samples were washed three times with PBS and mounted in Mounting Medium with DAPI (Solarbio). Finally, images were acquired using an Andor spinning disk or Nikon confocal microscopy system. All images were captured and analyzed in a blinded manner.

### Organotypic hippocampal slice culture, transfection and imaging

We prepared organotypic hippocampal slice cultures following previously established protocols (Chen et al., 2014). Briefly, slices were obtained from P7-8 Sprague-Dawley rats. At DIV 3, slices were biolistically transfected using a gene gun (Bio-Rad). Gold particles (1.6 μm in diameter; Bio-Rad) were coated with 25 μg of cDNA along with 5 μg of GFP. Live cell imaging was conducted at DIV9/10 using a Nikon confocal system equipped with a water-immersion 60×objective. Protrusions from dendrites longer than 0.4 μm were counted as spines. For spine density analysis, secondary basal dendrites were selected, and 2-5 different dendrites were imaged from each pyramidal cell. Both image acquisition and spine counting were carried out in a blinded manner.

### Aβ40 and Aβ42 ELISA

We analyzed all cell sample sets using Aβ40 and Aβ42 ELISA kits (Invitrogen, KHB3481 and KHB3441) according to the manufacturer’s instructions. The cells were sonicated in RIPA buffer (Solarbio) containing a protease inhibitor cocktail (Selleck), followed by centrifugation at 12,000 ×g for 10 minutes to prepare the samples.

### Lentivirus production

We inserted β-CTF-flag and GFP into the pFHTrePW vector (Ni et al., 2023) backbone under the control of the Tet Response Element (TRE 3G). The rtTA for the tet-off system was expressed under the control of the human synapsin I promoter. The pFHTrePW plasmid containing β-CTF-flag or GFP, along with PSPAX2 and PMD2g, was transfected into HEK293T cells. After 48 hours, the cell culture medium containing the virus was collected and filtered through a 0.45μm syringe filter (Millipore). Subsequently, it was concentrated using ultracentrifugation, aliquoted, and stored at -80 ℃.

### Adeno-Associated virus production

We performed AAV purification using triple-transfected HEK293T cells. Briefly, we transfected pHelper, pAAV2-8, and pAAV-MCS containing GFP or β-CTF into HEK293T cells at a ratio of 1:1:1. After 72 hours of incubation, the culture medium was collected in a 50 mL centrifuge tube, and the cell pellet was collected in another tube with lysis buffer (50 mM Tris-HCl, 150 mM NaCl, 2 mM MgCl_2_, pH 8.0). The cell pellet was subjected to three cycles of freezing and thawing, followed by mixed with the culture medium. PEG8000 (10%) and NaCl (1 M/L) were added to the mixture. After incubating for 1 hour on ice, the mixture was centrifuged for 15 minutes at 12,000 ×g and 4 °C. The precipitate was gently resuspended in lysis buffer containing DNase I (Roche) using a 1000-ml pipette. After a 30-minute incubation at 37 °C, chloroform in a 1:1 (v:v) ratio was added to each tube, shaken for 1 hour, and then centrifuged for 15 minutes at 12,000 ×g and 4°C. To harvest the concentrated AAV, the supernatant was processed using an ultrafiltration column (Millipore).

### Electrophysiology

We performed patch-clamp recordings from hippocampal pyramidal cells in acute brain slices. Adult mice approximately 8 weeks old were anesthetized with isoflurane and then decapitated. The hippocampus was harvested intact and placed into cold choline cutting buffer (110 mM Choline Cl, 2.5 mM KCl, 1.25 mM NaH_2_PO_4_, 25 mM NaHCO_3_, 7 mM MgSO_4_, 25 mM D-glucose, 3.1 mM Na Pyruvate, 11.6 mM Na Ascorbate, 0.5 mM CaCl_2_ at pH 7.25). Using a vibrating microtome (Leica VT1200S), the hippocampus was sliced into 400 μm sections, followed by a 30-minute recovery period at 34 ℃ and another 30-minute recovery period at room temperature in artificial cerebrospinal fluid (ACSF).

The recording external solution consisted of ACSF containing (in mM): 127 NaCl, 2.5 KCl, 12.5 NaH_2_PO_4_, 25 NaHCO_3_, 25 D-glucose, 2.5 CaCl_2_, and 1.3 MgCl_2_, aerated with 95% O_2_/5% CO_2_ to maintain a pH around 7.25. Pipettes (TW150F-4, World Precision Instruments) with a resistance of 4–6 mΩ were filled with an internal solution containing (in mM): 115 cesium methanesulfonate, 20 CsCl, 10 HEPES, 2.5 MgCl_2_, 4 Na_2_ATP, 0.4 Na_3_GTP, 10 Na phosphocreatine, and 0.6 EGTA (pH 7.25). Miniature excitatory postsynaptic currents (mEPSCs) were recorded by whole-cell voltage clamp. The mEPSCs were recorded at a holding potential of -70 mV in the presence of 1 μM tetrodotoxin (TTX, MedChem express, HY-12526A) and 100 μM picrotoxin (PTX, Sigma, P1675). EPSCs were analyzed using MiniAnalysis software (Synaptosoft) with an amplitude threshold of 5 pA.

### Antibodies and drugs

The following antibodies were used in this study: Y188 antibody (Abcam, ab32136), D10E5 antibody (Cell Signaling, 5606), Flag antibody (Sigma-Aldrich, F3165), β-Tubulin III antibody (Sigma, T 2200), GAPDH antibody (Proteintech, 60004-1-Ig), HA antibody (Cell Signaling, 3724S), GFP antibody (Abcam, ab13970), SYN antibody (Millipore, AB1543), GluR1 antibody (Millipore, MAB2263), GluR2 antibody (Millipore, AB1768-1). Peroxidase-Conjugated secondary antibodies: Goat Anti-Rabbit IgG (YESEN, 33101ES60), Goat Anti-Mouse IgG (YESEN, 33201ES60). Fluorescent secondary antibodies: Alexa 488 Goat anti-chicken IgG (Thermo Fisher, A11039), Alexa 568 Goat anti-chicken IgY (Thermo Fisher, A11041), Alexa 568 Goat anti-rabbit IgG (Thermo Fisher, A11036), Alexa 647 Goat anti-Mouse IgG (Thermo Fisher, A21236). Inhibitors: PF-03084014 (Selleck, S7731).

### Plasmids construction

We sub-cloned the expression of GFP, BACE1-GFP, GFP-Rab5, APP, APP-IRES-BACE1 or related mutations into the pCAGGS mammalian expression vector. The AAV constructs of GFP and β-CTF-HA were sub-cloned into the pAAV-MCS vector. Lentivirus constructs of GFP, β-CTF, and β-CTF-IRES-GFP were sub-cloned into the pFHTrePW vector backbone under the control of the TRE 3G.

### Behavioral Analysis

Animals underwent a 1-hour habituation period in the room before all behavioral tests. All tests were conducted and analyzed in a blinded manner.

### Open field test (OFT)

The mice were introduced into a 40 ×40 cm box without roofs and permitted to explore the apparatus freely for 1 hour. Their activities were monitored using the Ethovision video tracking system (Noldus Information Technology Inc., Leesburg, VA, USA). The field was divided into a center area (24 ×24 cm) and the whole arena. Parameters such as total travel distance, duration in the center area, velocity, and rearing frequency were automatically recorded and analyzed.

### Y maze

Each mouse was placed in one arm of a Y-maze (with arms measuring 30 cm in length) and allowed to explore the maze freely for 8 minutes. The movements of the mice were captured by a video camera positioned above the arena. The number of alternations and entries were analyzed using the Ethovision video tracking system.

### Elevated plus maze (EPM)

Anxiety-like behavior was assessed using an elevated plus maze (EPM), which is an elevated apparatus shaped like a plus sign (+). It consists of two open arms (35 ×5 × 0.3 cm), two closed arms (36 ×5 ×18 cm) with 15 cm walls and open roofs, and a 5 × 5 cm central area. At the beginning of the test, each mouse was placed in the center of the maze facing an open arm and allowed to explore the maze freely for 5 minutes. The distance explored and time spent in the apparatus were recorded and analyzed using the SMART software (Panlab, Barcelona, Spain).

### Water T maze (WTM)

Spatial learning and memory were evaluated using the Water T Maze (WTM) behavioral paradigm. In this task, mice were trained to utilize spatial cues within a room to locate a concealed platform and escape from water. In the reversal test, the hidden platform was relocated to the opposite arm to assess cognitive flexibility. The testing apparatus consisted of a plus maze, with each arm measuring 45 cm in length and 10 cm in width, constructed from clear Plexiglas. Each arm was designated as N, S, E, or W. A divider was positioned to block off an arm, allowing the mice to choose only the E or W arm for escape. The water temperature was maintained at 25-26 °C and made opaque by adding white, nontoxic powder. An escape platform was submerged 1 cm below the water’s surface on the E side of the maze, rendering it invisible to the mice.

At the beginning of each trial, the divider was put in place to block off the appropriate arm. Mice were placed at the starting point and allowed to explore the apparatus freely to reach the platform. If the mouse reached the platform within one minute, the response was considered correct; otherwise, it is considered incorrect. The experimenter recorded whether each response was correct or incorrect, and mice were permitted to remain on the platform for 10 seconds before being removed. Each day, mice underwent 5 trials with starting points semi-randomized between the N and S positions. The criterion for acquisition was achieving 80% or more correct responses averaged across the 5 trials for 2 consecutive days. After acquisition training, reversal training commenced. The hidden platform was relocated to the opposite side (W), and the same procedure was repeated until the mice successfully learned the new platform position. Mice that achieved 80% or more correct responses across the 5 trials for 2 consecutive days were deemed to have passed the test.

### Fear conditioning (FC)

The fear conditioning test comprised two components: contextual fear conditioning and cued fear conditioning. During the training stage, mice were introduced into conditioning boxes and allowed to freely explore the environment for 3 minutes. The baseline freezing behavior was recorded using a visual camera and analyzed with the Ethovision video tracking system. Subsequently, mice were exposed to an 80 dB, 2 kHz tone for 30 seconds (conditioned stimulus, CS), during which an inescapable 0.3 mA foot shock (unconditioned stimulus, US) was delivered in the last 2 seconds of the tone. This procedure was repeated three times with a 30-second inter-stimulus-interval.

For the contextual fear conditioning test, conducted on the second day, mice were placed in the same conditioning box used on the previous day for 5 minutes, and their freezing behavior was recorded for further analysis. In the cued fear conditioning test, conducted on the third day after training and contextual fear conditioning, mice were placed in a different chamber from the previous one. After a 3-minute free exploration period, mice were exposed to a 3-minute tone stimulus (2 kHz, 80 dB). Freezing behavior during the 3-minute tone stimulus was measured and analyzed using the Ethovision video tracking system.

### Tail suspension test (TST)

Each mouse was suspended by its tail using tape affixed to the ceiling of a three-walled rectangular compartment measuring 30 cm in height, 15 cm in width, and 15 cm in depth. The mice dangled downward, with ample space provided for movement. Video recording was conducted from the side, capturing the animals’ immobile time (defined as cessation of limb movements for more than 2 seconds) during the final 4 minutes of a 6-minute session.

### Stereotactic virus injection

Mice aged 4 weeks or 7-8 weeks received injections of lentivirus or AAV in the hippocampus at coordinates ML:±1.5 mm, AP: -2 mm, DV: -1.5 mm. For spine density analysis, mice were harvested after 2 months; for behavioral tests, mice were evaluated after 4 months.

### Liquid chromatography tandem mass spectrometry (LC-MS/MS) sample preparation and analysis

For protein sample preparation, neurons seeded in 24-well plates underwent lysis using ultrasonic waves in RIPA buffer. After centrifugation at 18,000 ×g for 10 minutes at 4 ℃, the supernatants were collected and quantified using a BCA assay (Thermo Fisher). Protein extracts were subjected to an in-solution digest protocol, involving reduction with 5 mM Tris(2-carboxyethyl) phosphine hydrochloride (TCEP) (Aldrich, USA) followed by alkylation with 10 mM iodoacetamide (IAA) (Sigma, USA). Trypsin was added at a 1:100 ratio, and the mixture was incubated at 37 ℃ overnight in the dark. Digested peptides were then collected by centrifugation and desalted using C18 tips (Pierce, USA) for subsequent analysis.

For LC-MS/MS analysis, the peptide mixture was analyzed using an online EASY-nLC 1000 HPLC system coupled with an Orbitrap Fusion mass spectrometer (Thermo Fisher Scientific). The sample was loaded directly onto a 15-cm homemade capillary column (100 μm I.D., C18-AQ 1.9 μm resin, Dr. Maisch). Mobile phase A consisted of 0.1% formic acid (FA), 2% acetonitrile (ACN), and 98% H_2_O, while mobile phase B comprised 0.1% FA, 2% H2O, and 98% ACN. A 180-minute gradient (mobile phase B: 2% at 0 min, 5% at 7 min, 20% at 127 min, 35% at 167 min, 95% at 173 min, 95% at 180 min) was employed at a static flow rate of 300 nl/min.

We acquired data for proteomic analysis in a data-dependent mode, starting with one full MS1 scan in the Orbitrap (m/z: 300-1800; resolution: 120,000; AGC target value: 500,000; maximal injection time: 50 ms), followed by an MS2 scan in the linear ion trap (32% normalized collision energy; maximal injection time: 250 ms). The isolation window was set at 1.6 m/z.

### Statistics

Data were processed using Microsoft Excel, and statistical analysis was performed with GraphPad Prism 7. Figures were made using Adobe Illustrator V26.3.1. Sample sizes and the statistical analyses performed are described in the respective figure legends. For all analyses, P<0.05 was considered statistically significant.

## Supporting information

Supplementary figures associated with main manuscript

## Acknowledgements

We thank the staff members of the Animal Facility at the National Facility for Protein Science in Shanghai (NFPS), Shanghai Advanced Research Institute, Chinese Academy of Sciences, China for providing assistance in mouse breeding and maintenance. This study was supported by Shanghai Basic Research Pioneer Project. This project received support from National Natural Science Foundation of China to Y.C. (31671044, 91849204); National Program on Key Research Project of China to Y.C. (2016YFA0501901); Shanghai Municipal Science and Technology Major Project to

Y.C. (Grant No. 2019SHZDZX02).

## Author contributions

Conceptualization: Yelin Chen, Yang Geng.; Methodology: Yelin Chen, Yang Geng., Mengxun Luo; Investigation and Validation: Mengxun Luo., Jia Zhou., Cailu Sun, Chaoying Fu, Wanjia Chen, Chenfang Si; Visualization: Mengxun Luo; Formal analysis: Mengxun Luo, Jia Zhou, Cailu Sun, Chenfang Si, Data curation: Mengxun Luo, Chenfang Si, Yaoyang Zhang; Writing – original draft preparation: Mengxun Luo; Writing – review & editing: Yelin Chen, Yang Geng; Supervision and Project administration: Yelin Chen; Funding acquisition: Yelin Chen.

## Conflict of Interest

The authors declare no competing interests.

## Notes

### Competing Interest Statement

The authors have declared no competing interest.

### Summary of Updates

Quantitative analyses of Figures 5A-C, 5I, 6J in Figures 5D, 5J and 6K. Figures 6E-G updated to clarify alterations in NMDA receptors and presynaptic SNARE complex led by β-CTF. Discussions revised. Supplementary figures updated.

## References

1. Abramov, E., Dolev, I., Fogel, H., Ciccotosto, G.D., Ruff, E., and Slutsky, I. (2009). Amyloid-beta as a positive endogenous regulator of release probability at hippocampal synapses. Nature neuroscience 12, 1567–1576.

2. Bittner, T., Fuhrmann, M., Burgold, S., Jung, C.K., Volbracht, C., Steiner, H., Mitteregger, G., Kretzschmar, H.A., Haass, C., and Herms, J. (2009). Gamma-secretase inhibition reduces spine density in vivo via an amyloid precursor protein-dependent pathway. The Journal of neuroscience : the official journal of the Society for Neuroscience 29, 10405–10409.

3. Blennow, K., de Leon, M.J., and Zetterberg, H. (2006). Alzheimer’s disease. Lancet (London, England) 368, 387–403.

4. Chapman, P.F., White, G.L., Jones, M.W., Cooper-Blacketer, D., Marshall, V.J., Irizarry, M., Younkin, L., Good, M.A., Bliss, T.V., Hyman, B.T., et al. (1999). Impaired synaptic plasticity and learning in aged amyloid precursor protein transgenic mice. Nature neuroscience 2, 271–276.

5. Chen, Y., Wang, Y., Ertürk, A., Kallop, D., Jiang, Z., Weimer, R.M., Kaminker, J., and Sheng, M. (2014). Activity-induced Nr4a1 regulates spine density and distribution pattern of excitatory synapses in pyramidal neurons. Neuron 83, 431–443.

6. Cheng, X., He, P., Lee, T., Yao, H., Li, R., and Shen, Y. (2014). High activities of BACE1 in brains with mild cognitive impairment. The American journal of pathology 184, 141–147.

7. Citron, M., Teplow, D.B., and Selkoe, D.J. (1995). Generation of amyloid beta protein from its precursor is sequence specific. Neuron 14, 661–670.

8. de Wilde, M.C., Overk, C.R., Sijben, J.W., and Masliah, E. (2016). Meta-analysis of synaptic pathology in Alzheimer’s disease reveals selective molecular vesicular machinery vulnerability. Alzheimer’s & dementia : the journal of the Alzheimer’s Association 12, 633–644.

9. DeKosky, S.T., Scheff, S.W., and Styren, S.D. (1996). Structural correlates of cognition in dementia: quantification and assessment of synapse change. Neurodegeneration : a journal for neurodegenerative disorders, neuroprotection, and neuroregeneration 5, 417–421.

10. Doody, R.S., Raman, R., Farlow, M., Iwatsubo, T., Vellas, B., Joffe, S., Kieburtz, K., He, F., Sun, X., Thomas, R.G., et al. (2013). A phase 3 trial of semagacestat for treatment of Alzheimer’s disease. The New England journal of medicine 369, 341–350.

11. Glenner, G.G., and Wong, C.W. (1984). Alzheimer’s disease: initial report of the purification and characterization of a novel cerebrovascular amyloid protein. Biochemical and biophysical research communications 120, 885–890.

12. Golde, T.E., Estus, S., Younkin, L.H., Selkoe, D.J., and Younkin, S.G. (1992). Processing of the amyloid protein precursor to potentially amyloidogenic derivatives. Science 255, 728–730.

13. Gómez-Isla, T., Price, J.L., McKeel, D.W., Jr., Morris, J.C., Growdon, J.H., and Hyman, B.T. (1996). Profound loss of layer II entorhinal cortex neurons occurs in very mild Alzheimer’s disease. The Journal of neuroscience : the official journal of the Society for Neuroscience 16, 4491–4500.

14. Güner, G., and Lichtenthaler, S.F. (2020). The substrate repertoire of γ-secretase/presenilin. Seminars in cell & developmental biology 105, 27–42.

15. Hu, X., Das, B., Hou, H., He, W., and Yan, R. (2018). BACE1 deletion in the adult mouse reverses preformed amyloid deposition and improves cognitive functions. The Journal of experimental medicine 215, 927–940.

16. Israel, M.A., Yuan, S.H., Bardy, C., Reyna, S.M., Mu, Y., Herrera, C., Hefferan, M.P., Van Gorp, S., Nazor, K.L., Boscolo, F.S., et al. (2012). Probing sporadic and familial Alzheimer’s disease using induced pluripotent stem cells. Nature 482, 216–220.

17. Jiang, Y., Mullaney, K.A., Peterhoff, C.M., Che, S., Schmidt, S.D., Boyer-Boiteau, A., Ginsberg, S.D., Cataldo, A.M., Mathews, P.M., and Nixon, R.A. (2010). Alzheimer’s-related endosome dysfunction in Down syndrome is Abeta-independent but requires APP and is reversed by BACE-1 inhibition. Proceedings of the National Academy of Sciences of the United States of America 107, 1630–1635.

18. JohnHardy, and Selkoe, D.J. (2002). The Amyloid Hypothesis of Alzheimer’s Disease: Progress and Problems on the Road to Therapeutics. Science 297, 353–356.

19. Jonsson, T., Atwal, J.K., Steinberg, S., Snaedal, J., Jonsson, P.V., Bjornsson, S., Stefansson, H., Sulem, P., Gudbjartsson, D., Maloney, J., et al. (2012). A mutation in APP protects against Alzheimer’s disease and age-related cognitive decline. Nature 488, 96–99.

20. Kamenetz, F., Tomita, T., Hsieh, H., Seabrook, G., Borchelt, D., Iwatsubo, T., Sisodia, S., and Malinow, R. (2003). APP processing and synaptic function. Neuron 37, 925–937.

21. Kessels, H.W., Nabavi, S., and Malinow, R. (2013). Metabotropic NMDA receptor function is required for β-amyloid-induced synaptic depression. Proceedings of the National Academy of Sciences of the United States of America 110, 4033–4038.

22. Kim, S., Sato, Y., Mohan, P.S., Peterhoff, C., Pensalfini, A., Rigoglioso, A., Jiang, Y., and Nixon, R.A. (2016). Evidence that the rab5 effector APPL1 mediates APP-βCTF-induced dysfunction of endosomes in Down syndrome and Alzheimer’s disease. Molecular psychiatry 21, 707–716.

23. Konietzko, U. (2012). AICD nuclear signaling and its possible contribution to Alzheimer’s disease. Current Alzheimer research 9, 200–216.

24. Kwart, D., Gregg, A., Scheckel, C., Murphy, E.A., Paquet, D., Duffield, M., Fak, J., Olsen, O., Darnell, R.B., and Tessier-Lavigne, M. (2019). A Large Panel of Isogenic APP and PSEN1 Mutant Human iPSC Neurons Reveals Shared Endosomal Abnormalities Mediated by APP β-CTFs, Not Aβ. Neuron 104, 256–270.e255.

25. Lai, A., Sisodia, S.S., and Trowbridge, I.S. (1995). Characterization of sorting signals in the beta-amyloid precursor protein cytoplasmic domain. The Journal of biological chemistry 270, 3565–3573.

26. Lee, S.E., Kwon, D., Shin, N., Kong, D., Kim, N.G., Kim, H.Y., Kim, M.J., Choi, S.W., and Kang, K.S. (2022). Accumulation of APP-CTF induces mitophagy dysfunction in the iNSCs model of Alzheimer’s disease. Cell death discovery 8, 1.

27. Leyns, C.E.G., and Holtzman, D.M. (2017). Glial contributions to neurodegeneration in tauopathies. Molecular neurodegeneration 12, 50.

28. Liu, C.Y., Ohki, Y., Tomita, T., Osawa, S., Reed, B.R., Jagust, W., Van Berlo, V., Jin, L.W., Chui, H.C., Coppola, G., et al. (2017). Two Novel Mutations in the First Transmembrane Domain of Presenilin1 Cause Young-Onset Alzheimer’s Disease. Journal of Alzheimer’s disease : JAD 58, 1035–1041.

29. Masters, C.L., Simms, G., Weinman, N.A., Multhaup, G., McDonald, B.L., and Beyreuther, K. (1985). Amyloid plaque core protein in Alzheimer disease and Down syndrome. Proceedings of the National Academy of Sciences of the United States of America 82, 4245–4249.

30. Mintun, M.A., Lo, A.C., Duggan Evans, C., Wessels, A.M., Ardayfio, P.A., Andersen, S.W., Shcherbinin, S., Sparks, J., Sims, J.R., Brys, M., et al. (2021). Donanemab in Early Alzheimer’s Disease. The New England journal of medicine 384, 1691–1704.

31. Mullan, M., Crawford, F., Axelman, K., Houlden, H., Lilius, L., Winblad, B., and Lannfelt, L. (1992). A pathogenic mutation for probable Alzheimer’s disease in the APP gene at the N-terminus of beta-amyloid. Nature genetics 1, 345–347.

32. Ni, J., Ren, Y., Su, T., Zhou, J., Fu, C., Lu, Y., Li, D., Zhao, J., Li, Y., Zhang, Y., et al. (2023). Loss of TDP-43 function underlies hippocampal and cortical synaptic deficits in TDP-43 proteinopathies. Molecular psychiatry 28, 931–945.

33. Nikolaev, A., McLaughlin, T., O’Leary, D.D., and Tessier-Lavigne, M. (2009). APP binds DR6 to trigger axon pruning and neuron death via distinct caspases. Nature 457, 981–989.

34. Nilsberth, C., Westlind-Danielsson, A., Eckman, C.B., Condron, M.M., Axelman, K., Forsell, C., Stenh, C., Luthman, J., Teplow, D.B., Younkin, S.G., et al. (2001). The ’Arctic’ APP mutation (E693G) causes Alzheimer’s disease by enhanced Abeta protofibril formation. Nature neuroscience 4, 887–893.

35. Oakley, H., Cole, S.L., Logan, S., Maus, E., Shao, P., Craft, J., Guillozet-Bongaarts, A., Ohno, M., Disterhoft, J., Van Eldik, L., et al. (2006). Intraneuronal beta-amyloid aggregates, neurodegeneration, and neuron loss in transgenic mice with five familial Alzheimer’s disease mutations: potential factors in amyloid plaque formation. The Journal of neuroscience : the official journal of the Society for Neuroscience 26, 10129–10140.

36. Oddo, S., Caccamo, A., Shepherd, J.D., Murphy, M.P., Golde, T.E., Kayed, R., Metherate, R., Mattson, M.P., Akbari, Y., and LaFerla, F.M. (2003). Triple-transgenic model of Alzheimer’s disease with plaques and tangles: intracellular Abeta and synaptic dysfunction. Neuron 39, 409–421.

37. Pensalfini, A., Kim, S., Subbanna, S., Bleiwas, C., Goulbourne, C.N., Stavrides, P.H., Jiang, Y., Lee, J.H., Darji, S., Pawlik, M., et al. (2020). Endosomal Dysfunction Induced by Directly Overactivating Rab5 Recapitulates Prodromal and Neurodegenerative Features of Alzheimer’s Disease. Cell reports 33, 108420.

38. Peters, F., Salihoglu, H., Rodrigues, E., Herzog, E., Blume, T., Filser, S., Dorostkar, M., Shimshek, D.R., Brose, N., Neumann, U., et al. (2018). BACE1 inhibition more effectively suppresses initiation than progression of β-amyloid pathology. Acta neuropathologica 135, 695–710.

39. Pleen, J., and Townley, R. (2022). Alzheimer’s disease clinical trial update 2019-2021. Journal of neurology 269, 1038–1051.

40. Puzzo, D., Privitera, L., Leznik, E., Fà, M., Staniszewski, A., Palmeri, A., and Arancio, O. (2008). Picomolar amyloid-beta positively modulates synaptic plasticity and memory in hippocampus. The Journal of neuroscience : the official journal of the Society for Neuroscience 28, 14537–14545.

41. Reinders, N.R., Pao, Y., Renner, M.C., da Silva-Matos, C.M., Lodder, T.R., Malinow, R., and Kessels, H.W. (2016). Amyloid-β effects on synapses and memory require AMPA receptor subunit GluA3. Proceedings of the National Academy of Sciences of the United States of America 113, E6526–e6534.

42. Salehi, A., Delcroix, J.D., Belichenko, P.V., Zhan, K., Wu, C., Valletta, J.S., Takimoto-Kimura, R., Kleschevnikov, A.M., Sambamurti, K., Chung, P.P., et al. (2006). Increased App expression in a mouse model of Down’s syndrome disrupts NGF transport and causes cholinergic neuron degeneration. Neuron 51, 29–42.

43. Selkoe, D.J. (2002). Alzheimer’s disease is a synaptic failure. Science (New York, NY) 298, 789–791.

44. Sims, J.R., Zimmer, J.A., Evans, C.D., Lu, M., Ardayfio, P., Sparks, J., Wessels, A.M., Shcherbinin, S., Wang, H., Monkul Nery, E.S., et al. (2023). Donanemab in Early Symptomatic Alzheimer Disease: The TRAILBLAZER-ALZ 2 Randomized Clinical Trial. Jama 330, 512–527.

45. Sperling, R.A., Donohue, M.C., Raman, R., Rafii, M.S., Johnson, K., Masters, C.L., van Dyck, C.H., Iwatsubo, T., Marshall, G.A., Yaari, R., et al. (2023). Trial of Solanezumab in Preclinical Alzheimer’s Disease. The New England journal of medicine 389, 1096–1107.

46. Tamayev, R., Matsuda, S., Arancio, O., and D’Adamio, L. (2012). β-but not γ-secretase proteolysis of APP causes synaptic and memory deficits in a mouse model of dementia. EMBO molecular medicine 4, 171–179.

47. Terry, R.D., Masliah, E., Salmon, D.P., Butters, N., DeTeresa, R., Hill, R., Hansen, L.A., and Katzman, R. (1991). Physical basis of cognitive alterations in Alzheimer’s disease: synapse loss is the major correlate of cognitive impairment. Annals of neurology 30, 572–580.

48. Thinakaran, G., and Koo, E.H. (2008). Amyloid precursor protein trafficking, processing, and function. The Journal of biological chemistry 283, 29615–29619.

49. Tyan, S.H., Shih, A.Y., Walsh, J.J., Maruyama, H., Sarsoza, F., Ku, L., Eggert, S., Hof, P.R., Koo, E.H., and Dickstein, D.L. (2012). Amyloid precursor protein (APP) regulates synaptic structure and function. Molecular and cellular neurosciences 51, 43–52.

50. Uhlén, M., Fagerberg, L., Hallström, B.M., Lindskog, C., Oksvold, P., Mardinoglu, A., Sivertsson, Å., Kampf, C., Sjöstedt, E., Asplund, A., et al. (2015). Proteomics. Tissue-based map of the human proteome. Science 347, 1260419.

51. Vaillant-Beuchot, L., Mary, A., Pardossi-Piquard, R., Bourgeois, A., Lauritzen, I., Eysert, F., Kinoshita, P.F., Cazareth, J., Badot, C., Fragaki, K., et al. (2021). Accumulation of amyloid precursor protein C-terminal fragments triggers mitochondrial structure, function, and mitophagy defects in Alzheimer’s disease models and human brains. Acta neuropathologica 141, 39–65.

52. van Dyck, C.H., Swanson, C.J., Aisen, P., Bateman, R.J., Chen, C., Gee, M., Kanekiyo, M., Li, D., Reyderman, L., Cohen, S., et al. (2023). Lecanemab in Early Alzheimer’s Disease. The New England journal of medicine 388, 9–21.

53. Vassar, R., Bennett, B.D., Babu-Khan, S., Kahn, S., Mendiaz, E.A., Denis, P., Teplow, D.B., Ross, S., Amarante, P., Loeloff, R., et al. (1999). Beta-secretase cleavage of Alzheimer’s amyloid precursor protein by the transmembrane aspartic protease BACE. Science 286, 735–741.

54. Vohra, B.P., Sasaki, Y., Miller, B.R., Chang, J., DiAntonio, A., and Milbrandt, J. (2010). Amyloid precursor protein cleavage-dependent and -independent axonal degeneration programs share a common nicotinamide mononucleotide adenylyltransferase 1-sensitive pathway. The Journal of neuroscience : the official journal of the Society for Neuroscience 30, 13729–13738.

55. Wang, X., Wang, Z., Chen, Y., Huang, X., Hu, Y., Zhang, R., Ho, M.S., and Xue, L. (2014). FoxO mediates APP-induced AICD-dependent cell death. Cell death & disease 5, e1233.

56. Wang, Z., Jackson, R.J., Hong, W., Taylor, W.M., Corbett, G.T., Moreno, A., Liu, W., Li, S., Frosch, M.P., Slutsky, I., et al. (2017). Human Brain-Derived Aβ Oligomers Bind to Synapses and Disrupt Synaptic Activity in a Manner That Requires APP. The Journal of neuroscience : the official journal of the Society for Neuroscience 37, 11947–11966.

57. Wei, W., Nguyen, L.N., Kessels, H.W., Hagiwara, H., Sisodia, S., and Malinow, R. (2010). Amyloid beta from axons and dendrites reduces local spine number and plasticity. Nature neuroscience 13, 190–196.

58. Willem, M., Tahirovic, S., Busche, M.A., Ovsepian, S.V., Chafai, M., Kootar, S., Hornburg, D., Evans, L.D., Moore, S., Daria, A., et al. (2015). η-Secretase processing of APP inhibits neuronal activity in the hippocampus. Nature 526, 443–447.

59. Xiao, C., Liu, Y., Xu, J., Gan, X., and Xiao, Z. (2018). Septal and Hippocampal Neurons Contribute to Auditory Relay and Fear Conditioning. Frontiers in cellular neuroscience 12, 102.

60. Xu, W., Weissmiller, A.M., White, J.A., 2nd, Fang, F., Wang, X., Wu, Y., Pearn, M.L., Zhao, X., Sawa, M., Chen, S., et al. (2016). Amyloid precursor protein-mediated endocytic pathway disruption induces axonal dysfunction and neurodegeneration. The Journal of clinical investigation 126, 1815–1833.

61. Zhang, X., and Song, W. (2013). The role of APP and BACE1 trafficking in APP processing and amyloid-β generation. Alzheimer’s research & therapy 5, 46.

62. Zhang, Y.W., Thompson, R., Zhang, H., and Xu, H. (2011). APP processing in Alzheimer’s disease. Molecular brain 4, 3.

63. Zhou, B., Lu, J.G., Siddu, A., Wernig, M., and Südhof, T.C. (2022). Synaptogenic effect of APP-Swedish mutation in familial Alzheimer’s disease. Science translational medicine 14, eabn9380.

64. Zhou, Y., Zhou, B., Pache, L., Chang, M., Khodabakhshi, A.H., Tanaseichuk, O., Benner, C., and Chanda, S.K. (2019). Metascape provides a biologist-oriented resource for the analysis of systems-level datasets. Nature communications 10, 1523.

