## Supplementary figures associated with main manuscript for "APP β-CTF triggers cell-autonomous synaptic toxicity independent of Aβ"

Supplementary figure 1. Different APP and BACE1 expression ratios impacted spine density differently.

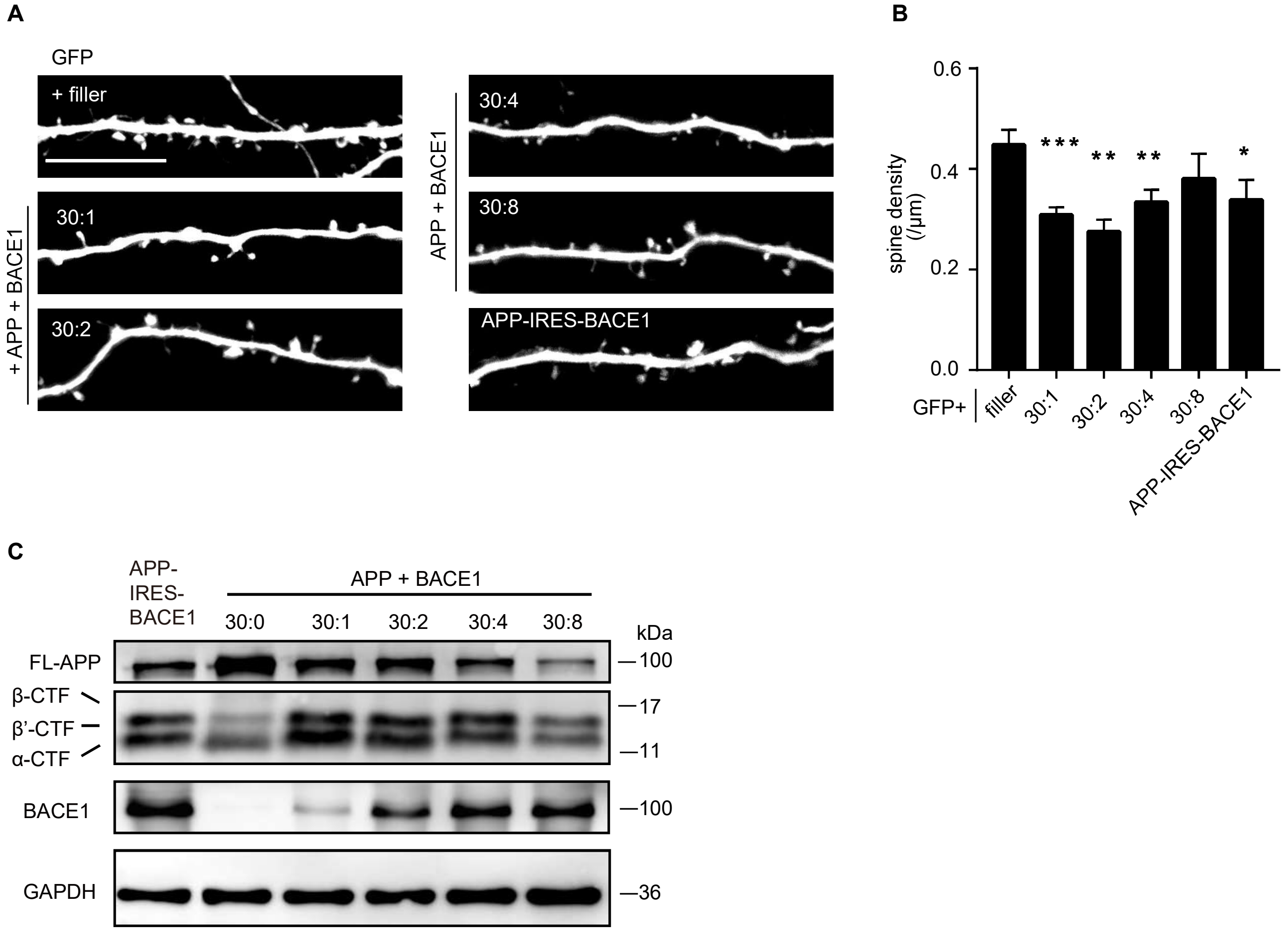

**Supplementary figure 1. Different APP and BACE1 expression ratios impacted spine density differently.**

**A-B** Representative images and measurements of the spine density from CA1 pyramidal neurons transiently expressing GFP or together with APP and BACE1 at different molecular ratios for 6-7 days in rat organotypic hippocampal slice cultures. (filler, n=8; 30:1, n=12; 30:2, n=5; 30:4, n=9; 30:8, n=6; APP-IRES-BACE1, n=9)

**C** Western blot of APP fragments (Y188) and BACE1 from HEK293T cells expressing APP and BACE1 at different ratios or APP-IRES-BACE1.

Scale bar, 10 μm. Statistics: One-way ANOVA. * $p < 0.05$, ** $p < 0.01$, *** $p < 0.001$, **** $p < 0.0001$. Error bars show SEM.

Supplementary figure 2. sAPP or AICD expression and γ-secretase inhibition did not affect dendritic spines.

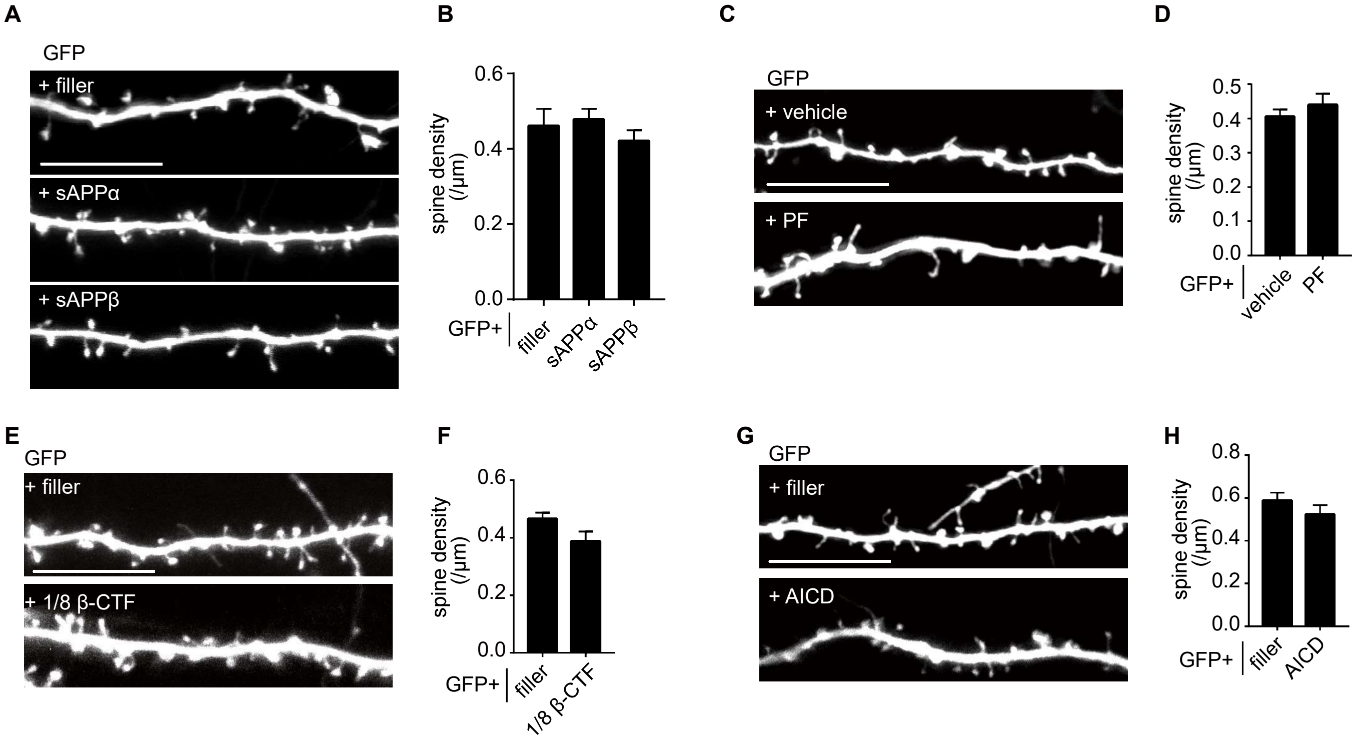

**Supplementary figure 2. sAPP or AICD expression and γ-secretase inhibition did not affect dendritic spines.**

**A-B** Representative images and spine density of basal dendrites from CA1 pyramidal neurons transiently expressing GFP alone or together with sAPPα or sAPPβ. (filler, n=10; sAPPα, n=13; sAPPβ, n=19)

**C-D** Representative images and spine density of basal dendrites from CA1 pyramidal neurons transiently expressing GFP treatment with vehicle or PF. (vehicle, n=13; PF, n=9)

**E-F** Representative images and spine density of basal dendrites from CA1 pyramidal neurons transiently expressing GFP alone or together with β-CTF (1/8 of the original plasmid amount). (filler, n=10; 1/8 β-CTF, n=11)

**G-H** Representative images and spine density of basal dendrites from CA1 pyramidal neurons transiently expressing GFP alone or together with AICD. (filler, n=6; AICD, n=10)

All dendritic images were acquired from rat organotypic hippocampal slice cultures after transfection for 6-7 days. PF, PF03084014, a γ secretase inhibitor. Statistics: One-way ANOVA or Student's t test. * $p < 0.05$, ** $p < 0.01$, *** $p < 0.001$, **** $p < 0.0001$. Error bars show SEM. Scale bars, 10 μm.

Supplementary figure 3. Additional neurobehavioral tests of mice expressing β-CTF in their brains.

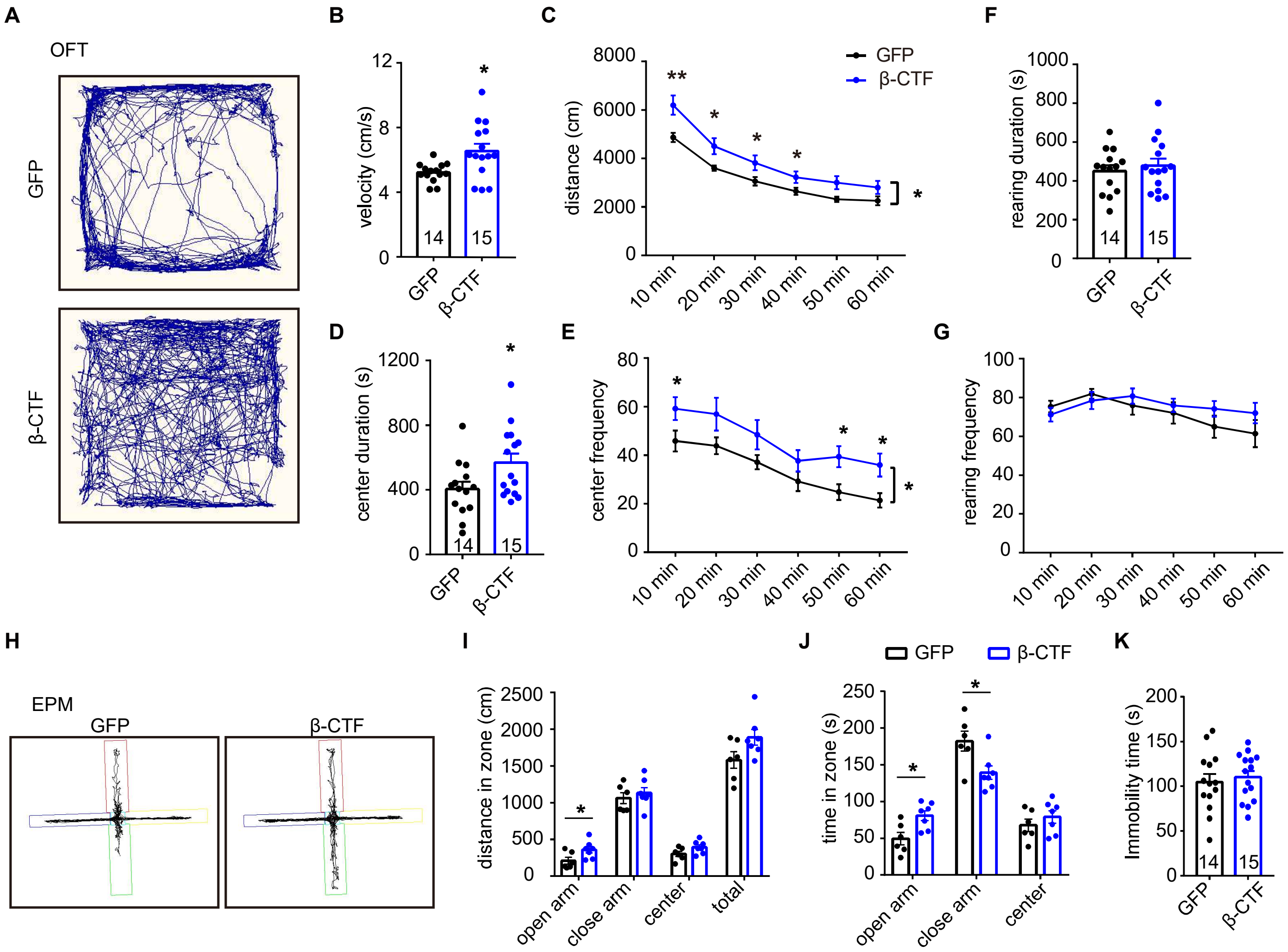

**Supplementary figure 3. Additional neurobehavioral tests of mice expressing β-CTF in their brains.**

**A** Representative travel traces of the open field tests from adult mice infected with AAV encoding GFP or β-CTF in their hippocampi.

**B-G** Quantitation of open field tests from adult mice infected with AAV encoding GFP or β-CTF in hippocampus. GFP, n=14; β-CTF, n=15.

**H** Representative traces of elevated plus maze tests from the mice infected with AAV encoding GFP or β-CTF in their hippocampi.

**I** Quantitation of elevated plus maze tests. GFP, n=6; β-CTF, n=7.

**J** Analysis of time spent in different arms of elevated plus maze. GFP, n=6; β-CTF, n=7.

**K** Immobility time from tail suspension tests. GFP, n=14; β-CTF, n=15.

Statistics: Repeated measures two-way ANOVA or Student's t test. * $p < 0.05$, ** $p < 0.01$, *** $p < 0.001$, **** $p < 0.0001$. Error bars show SEM.

Supplementary figure 4. Co-immunostaining of APP CTFs and a lysosomal marker.

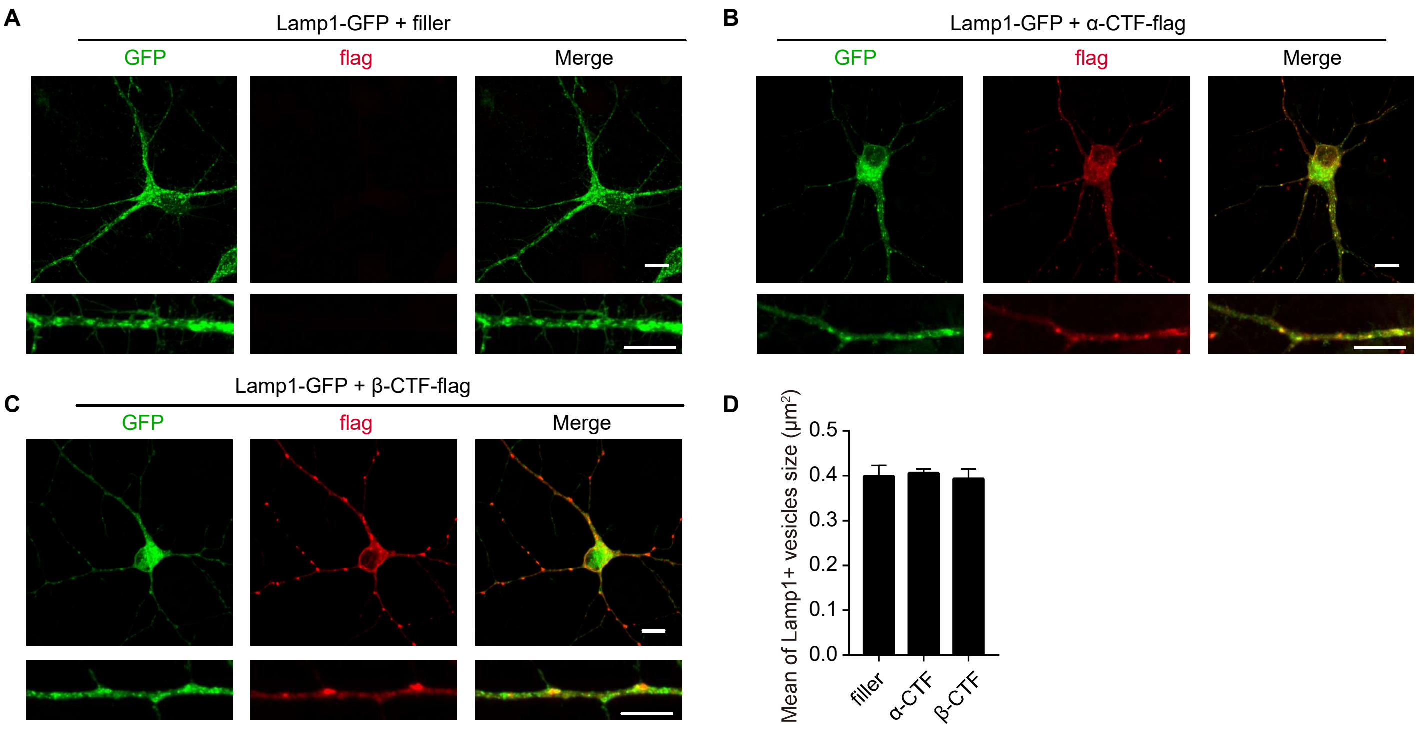

**Supplementary figure 4. Co-immunostaining of APP CTFs and a lysosomal marker.**

**A** Immunofluorescence staining of GFP (green) from dissociated rat hippocampal neurons expressing Lamp1-GFP.

**B** Immunofluorescence staining of GFP (green) and flag (red) from neurons expressing Lamp1-GFP and α-CTF-flag.

**C** Immunofluorescence staining of GFP (green) and flag (red) from neurons expressing Lamp1-GFP and β-CTF-flag.

**D** Quantitation of average Lamp1+ vesicles size of neurons expressing Lamp1-GFP only or co-expressing α/β-CTF-flag and Lamp1-GFP. (filler, n=6; α-CTF, n=7; β-CTF, n=7)

Scale bar, 10 μm. Statistics: One-way ANOVA. * $p < 0.05$, ** $p < 0.01$, *** $p < 0.001$, **** $p < 0.0001$. Error bars show SEM.

Supplementary figure 5. GO and Reactome Gene sets analysis of β-CTF-upregulated proteins in neurons.

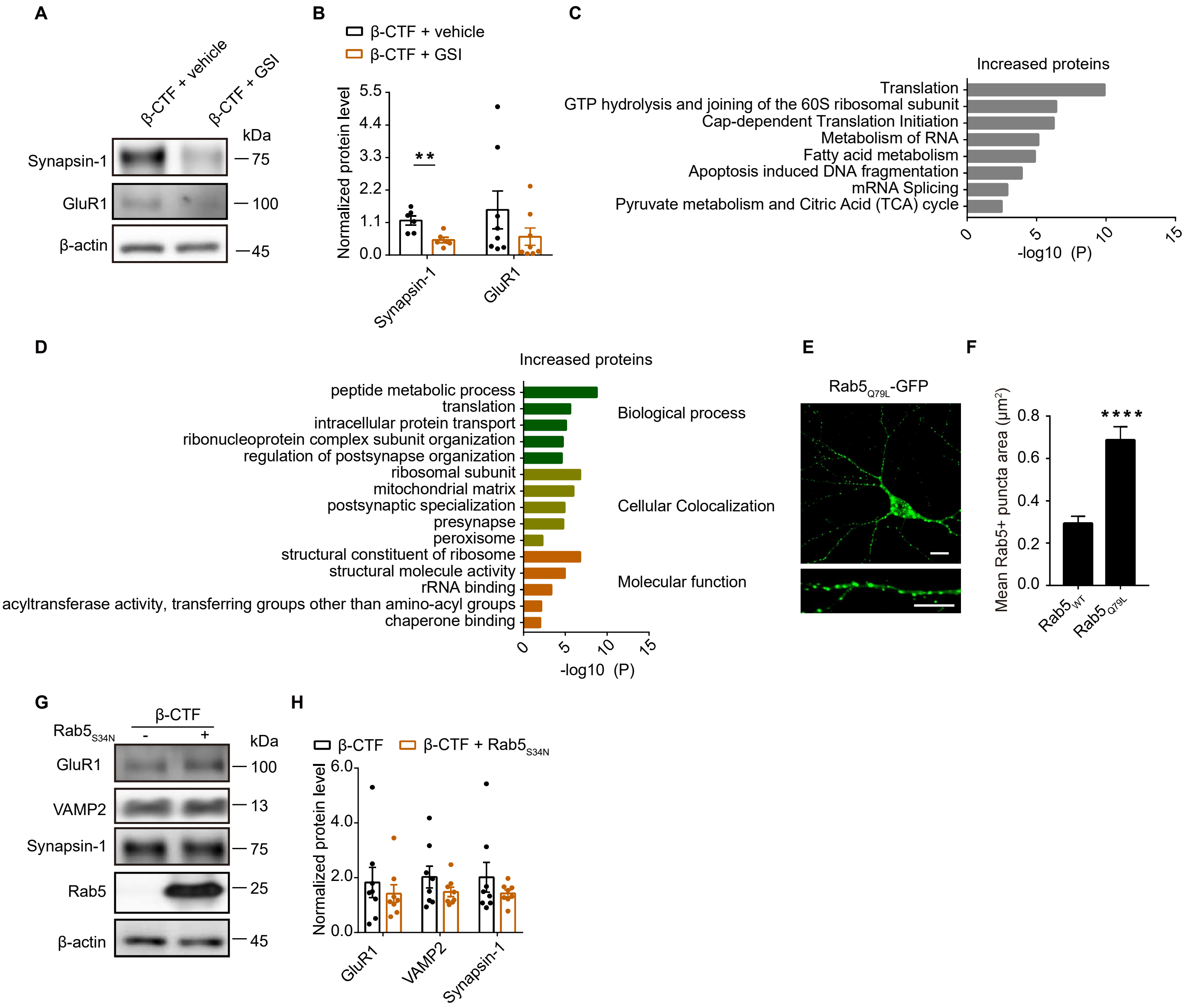

**Supplementary figure 5. GO and Reactome Gene sets analysis of β-CTF-upregulated proteins in neurons.**

**A-B** Western blot and corresponding statistic analysis of Synapsin-1 and GluR1 in dissociated hippocampal neurons infected with lentivirus expressing β-CTF treated with vehicle or PF. β-actin was used as an internal control.

**C** The Gene Ontology (GO) processes analysis of increased protein by Metascape.

**D** The Reactome Gene Sets processes analysis of increased protein by Metascape.

**E** Immunofluorescence staining of GFP (green) from dissociated rat hippocampal neurons expressing $Rab5_{Q79L}$-GFP.

**F** Quantitation of average Rab5+ puncta size from dissociated rat hippocampal neurons expressing $Rab5_{WT}$-GFP or $Rab5_{Q79L}$-GFP. ($Rab5_{WT}$, n=10; $Rab5_{Q79L}$, n=5)

**G-H** Western blot and corresponding statistic analysis of GluR1, VAMP2 and Synapsin-1 in dissociated hippocampal neurons infected with lentivirus expressing β-CTF or co-expressing β-CTF and $Rab5_{S34N}$. β-actin was used as an internal control.

Statistics: Student's t test. * $p < 0.05$, ** $p < 0.01$, *** $p < 0.001$, **** $p < 0.0001$. Error bars show SEM. Scale bar, 10 μm.
